# A Quantitative Proteome Map of the Human Body

**DOI:** 10.1101/797373

**Authors:** Lihua Jiang, Meng Wang, Shin Lin, Ruiqi Jian, Xiao Li, Joanne Chan, Huaying Fang, Guanlan Dong, GTEx Consortium, Hua Tang, Michael P. Snyder

## Abstract

Determining protein levels in each tissue and how they compare with RNA levels is important for understanding human biology and disease as well as regulatory processes that control protein levels. We quantified the relative protein levels from 12,627 genes across 32 normal human tissue types prepared by the GTEx project. Known and new tissue specific or enriched proteins (5,499) were identified and compared to transcriptome data. Many ubiquitous transcripts are found to encode highly tissue specific proteins. Discordance in the sites of RNA expression and protein detection also revealed potential sites of synthesis and action of protein signaling molecules. Overall, these results provide an extraordinary resource, and demonstrate that understanding protein levels can provide insights into metabolism, regulation, secretome, and human diseases.

**Summary:** Quantitative proteome study of 32 human tissues and integrated analysis with transcriptome data revealed that understanding protein levels could provide in-depth knowledge to post transcriptional or translational regulations, human metabolism, secretome, and diseases.

## Introduction

Understanding which components are expressed in which tissues is fundamental for studying human biology and disease. To date most efforts have focused on RNA because it is relatively easy to quantify. However, RNA expression does not directly impact phenotype as many additional levels of regulation including post-transcriptional, translational, and protein modifications all contribute to individual traits. Proteins, which reside downstream of transcription and participate in vital activities of cells, are ideal molecular phenotypes for understanding the contribution of post-transcriptional regulatory mechanisms to organismic level complex phenotypes. Previous studies have indicated that protein levels correlate poorly with transcript levels (*1*–*3*). Therefore, a detailed description of protein expression across tissues may provide complementary information to transcriptomic studies and inform how protein levels relate to human biology and disease.

Both mass spectrometry and immunolocalization studies have been performed to generate tissue maps of protein expression (*4, 5*). Mass spectrometry analysis of various cell lines and human tissues has identified approximately 85% of the proteins encoded by the 20,000 human protein coding genes, and provides an excellent first generation tissue map (*6*–*8*). The Human Protein Atlas project (HPA) generated a tissue-based map of the human proteome based on immunolocalization data across 32 tissues and 44 cell lines (*4*). However, the quantitative information in HPA primarily relied on transcriptome data. Overall, a quantitative comparison of protein expression across tissues, a direct assessment of correlation between protein and RNA in the same tissue samples, and the relationship of protein expression to biological processes (e.g. metabolism, secretion), are lacking.

The Genotype-Tissue Expression (GTEx) project provides unique opportunities to fill these gaps. GTEx collected samples from 54 tissues of 948 post-mortem donors, all of which have been transcriptomically characterized by RNA-seq (*9*–*13*). We quantified the relative levels of proteins from 12,627 genes across 32 normal human tissues in 201 tissue samples collected by the GTEx project. 5,499 known and novel tissue enriched and specific proteins were identified. Notably, many RNA transcripts do not always show concordant tissue enriched or specific patterns with their encoded proteins. As an example, many vesicular transport proteins involved in neurotransmitter function and cell-cell signaling are highly enriched in the brain but do not exhibit RNA enrichment. Conversely, tissue enriched/specific RNA transcripts can be found in tissues with no corresponding protein enrichment. We used the protein /RNA concordant enrichment information to suggest proteins that undergo constitutive and regulated secretion and the possible targeting locations of many secreted proteins. Examination of many tissue specific proteins reveals that they are associated with specific diseases and provide a molecular explanation for the underlying defects. Overall, these results provide a valuable resource, and demonstrate that understanding protein levels can provide insights into metabolism, regulation, sites of function, and human disease.

## Results

### Protein Profiling Across Tissues

We quantitatively profiled the proteome of 201 samples from 32 different tissue types of 14 normal individuals as shown in Figure. 1A. The samples covered all major organs (Table S1). The proteome data were acquired using a Tandem Mass Tag (TMT) 10plex/MS3 mass spectrometry strategy using a Fusion Orbitrap (Fig. 1B). TMT10plex enables ten isotopically labeled samples to be analyzed in a single experiment, which increases throughput and reduces technical variation (*14*). A common reference, created by mixing peptide digests from all 201 samples was included in each run. One technical challenge is the broad dynamic range of proteomes across different tissues. For example, in heart and muscle, a small number of top proteins can account for more than half of the mass of the entire proteome (*5, 15*). The high signals from these abundant proteins can suppress the signals from the lower abundance proteins leading to inaccurate quantitation unless extensive fractionation of each sample is carried out. To minimize this problem, we randomized the tissue samples such that each TMT10-plex consists of a mixture of tissues, and each multiplexed sample was extensively fractionated (Fig. 1B).

**Fig. 1.**
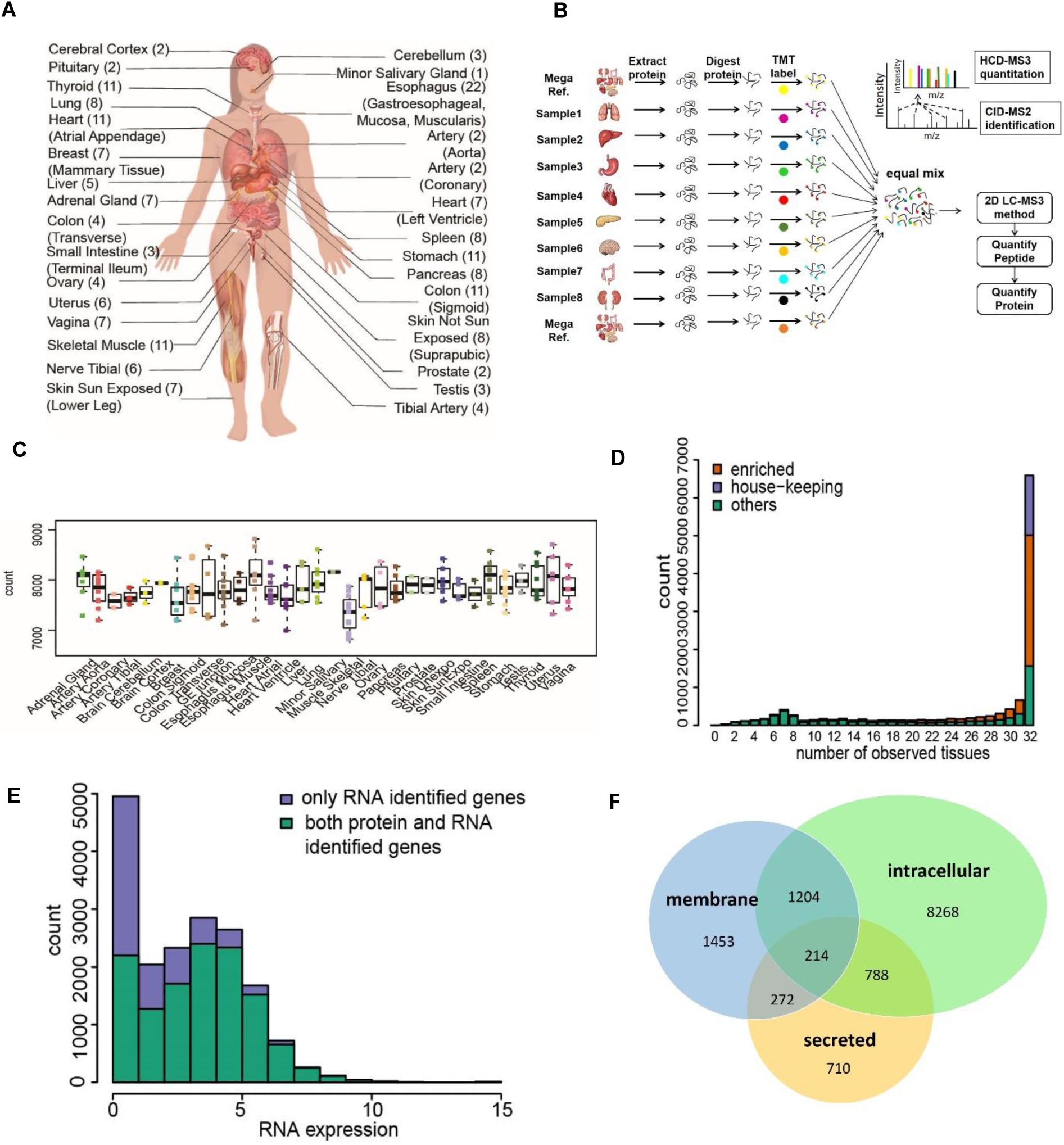
Overview of tissue proteome workflow and results. A. Type of tissues and biological replicates analyzed in this study. B. TMT 10plex and MS3 based mass spectrometry quantitative proteomics workflow. C. Number of proteinsquantitatedin each tissue. Each dot represents data from one person. D. Distribution of the number of proteins quantified across different numbers of tissues. The enrichment category is defined in Suppl Sec 3.1-3.2.E. Distribution of RNA TPM expression in log scale. The RNA with TPM less than 1 is collapsed to 1 here. The chi-square test is applied to test whether protein identification is independent to RNA expression (Supp Sec 4 and Table S8(a)).F. Category of proteins and numbers identified in this study. The predicated protein classes are from the results of HPA.

In total, we identified proteins encoded by 13,813 genes at peptide FDR of 1% for each sample and 12,627 were quantitated after applying strict filters (Table S1). For each tissue type, an average of more than 7,500 proteins were quantified (Fig. 1C) and there are 6,357 proteins present in all 32 tissue types (Fig. 1D). This result indicates that there is a core set of ubiquitously expressed proteins, consistent with previous studies, which have shown that an individual tissue is not strongly characterized by the simple presence or absence of proteins but rather by quantitative differences (*15*). To obtain a confident list of the total number of proteins detected across all tissues, we pooled all the mass spectra and performed a single database search. In total, 10,312 proteins were identified at protein FDR 1%. A summary of detected proteins is presented in Table S1 along with their confidence level of detection.

To determine classes of proteins that might be missing, we overlapped the corresponding RNA abundance of all the proteins identified in our study with the entire transcriptome (Fig. 1E). The results shown that there is much less protein identification when the corresponding RNA TPM is low. However, when RNA log2TPM is above 5, protein detection is not significantly affected by RNA abundance (Table S2&8). Based on the genome annotation, there are 5500 proteins predicted to be membrane-bound and 3000 secreted (*4*). We detected 3143 membrane proteins and 1984 secreted proteins (Fig. 1F); the membrane proteins were under detected across the entire RNA expression abundance level, presumably due to sample preparation methods. Among all the undetected proteins, a significant portion are expressed as RNA in the testis. Undetected proteins may be due to expression in a limited number of cell types, post-transcriptional regulation, degradation of proteins, or protein secretion, which is discussed further below.

### Tissue Enriched/Specific Proteins

To quantify the relative expression of each protein across tissues we developed and applied an in-house normalization algorithm. Compared to existing methods such as probability quotient normalization (*16*), this method identifies proteins separated from the population distribution and also enables more accurate protein quantification (*17*). After data normalization, samples were clustered based on their protein expression levels using hierarchical clustering based on pairwise Euclidean distance from Ward’s method (Fig S2). Samples were clearly separated by tissue types (Fig. 2A&S2) indicating that the differences among different tissue types exceed that of individuals. Physiologically related samples also clustered together. For example, arteries from different parts of the body were tightly clustered, as were heart and skeletal muscles. Although anatomically the stomach, small intestine, transverse colon and sigmoid colon are all part of the digestive system, the sigmoid colon samples did not cluster with the other members. This is likely because the sigmoid colon samples are largely composed of muscularis layer, and thus they cluster with other muscularis tissue samples, such as esophagus muscularis and esophagus-gastric junction, even though the esophagus is anatomically further away. Likewise, samples from esophagus mucosa layers were clustered with skin instead of other esophagus, likely due to functional similarity between epithelial cells. Interestingly lung was found to be tightly clustered with spleen despite their distinct functions. Further analysis revealed a common group of 78 proteins involved in immunity were enriched in both lung and spleen. Recent studies have shown that lung hosts many immune cells and is a large reservoir of neutrophils, likely accounting for the similarity (*18*).

**Fig. 2.**
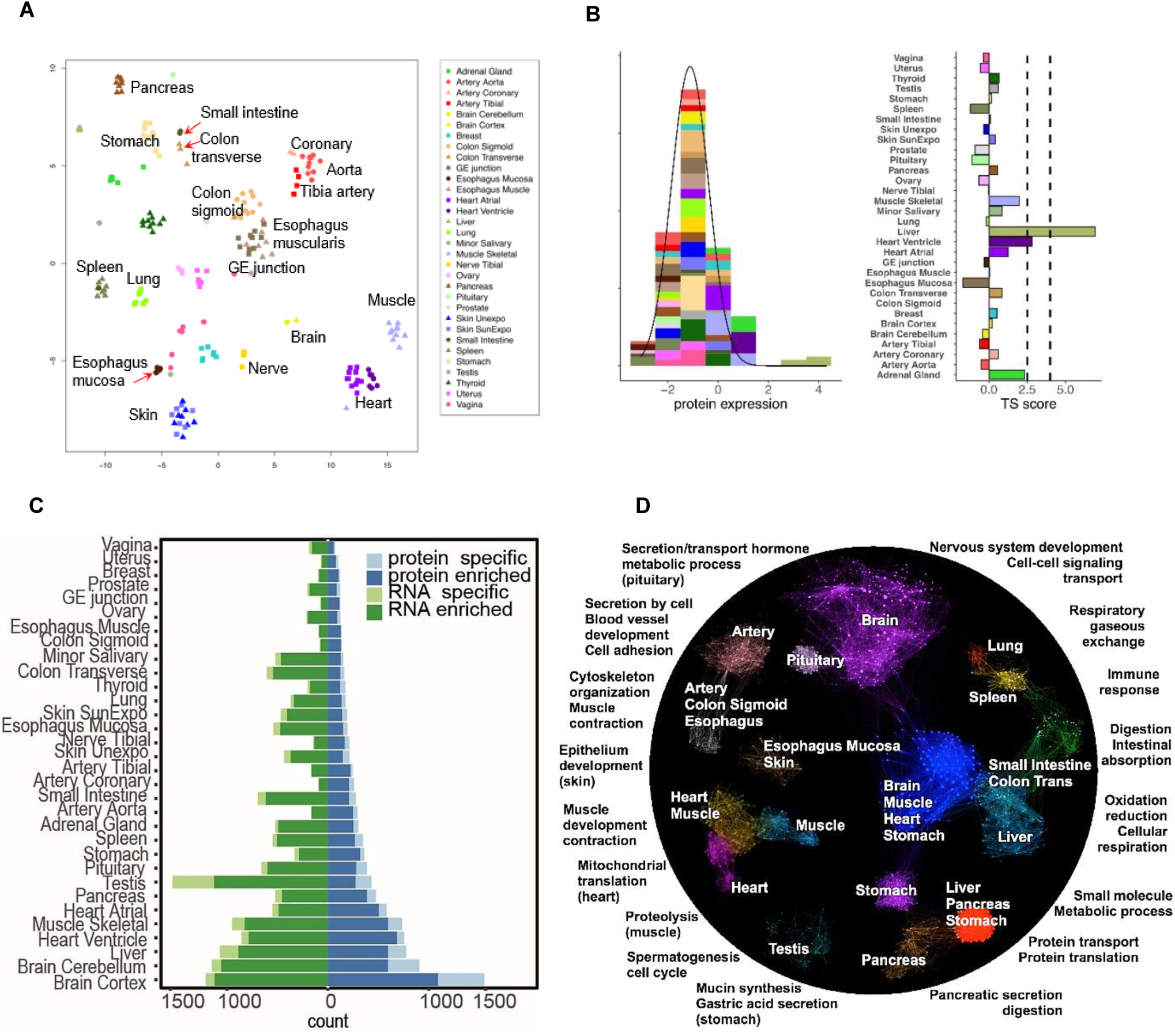
Quantitative proteome analysis across tissues. A. Clustering of proteome data across tissues from t SNE. Tissue type is the major factor that c lusters separates the samples from the same/different tissues. B. Method for defining Tissue Specificity (TS) scores. As an example, for gene PHYH, the left panel shows the distribution of its TS scores across tissues fitted using AdaTiSS (Supp Sec 3.1-3.2). The right panel shows its TS scores in each tissue. The vertical lines are at 2.5 and 4. C. The number of enriched and specific proteins/RNA across tissues. The enrichment categories are defined in Suppl Sec 3.1-3.2. D. Protein enrichment across tissues and their biological functions. The enriched proteins represent tissue specific/shared functions. The GO term functional enrichment results are summarized in Table S3.

The enrichment information of each protein across tissues was defined by Tissue Specificity (TS) score. The TS-score is defined as the robust estimation of the distance, measured in units of standard error, between median abundance in a tissue and the population mean. Figure. 2B showed the distribution of TS-scores for protein *PHYH* across all tissues. We consider a TS-score greater than 2.5 as an outlier, and define a protein to be tissue enriched if its TS-score reaches 2.5 in at least one tissue. If the TS-score of a protein is greater than 4 in a tissue and is at least 1.5 standard deviations away from the protein’s TS-scores in any other tissue, this protein is considered tissue specific. In total, there were 3,851 (31.5%) enriched proteins and 1,558 (12.7%) tissue specific proteins (Table S3&6). Brain has the highest number of enriched and specific proteins followed by liver, heart and muscle (Fig. 2C, Table S3).

Figure. 2D showed the enriched/specific proteins across tissues. As shown in the figure, some proteins are only enriched in one tissue and some are enriched in multiple tissues. GO analysis reveals that proteins only enriched in one tissue have functions that are highly tissue-specific, and proteins enriched in more than one tissue usually present shared functions (Table S4). For example, proteins involved in nervous system development and synaptic transmission are highly enriched in brain and a fraction of them is also enriched in the pituitary. However, in the pituitary, a group of peptide hormones are specifically enriched representing highly specialized functions of pituitary. Sarcomeric proteins which are essential for muscle development and contraction are highly enriched in both heart and skeletal muscle. The tissue specificities were further defined through the differential enrichment of multiple isoforms of the major sarcomeric proteins such as myosin, tropomyosin and troponin (Table S10). In addition, proteins involved in mitochondrial translation are more enriched in heart whereas proteins in proteolysis are more enriched in skeletal muscle. Interestingly, the left heart ventricle and heart atria also show different protein enrichments. Proteins involved in energy production are more enriched in left heart ventricle but peptide hormones (*NPPA, NPPB*) and specific myosin isoforms are only enriched in atria (Table S10). Proteins involved in oxidation and reduction are enriched in multiple metabolically active tissues such as heart, muscle, brain, liver and stomach, which will be further discussed in the metabolism section. Although ribosomal proteins are present in almost all organs, they are highly enriched in pancreas followed by liver and stomach; these organs are highly active in protein synthesis, especially the pancreas. Lung, spleen and small intestine share a group of proteins involved in immune response, but spleen has many more immune related proteins.

Proteins which are present in all tissues and not enriched in any of them are defined as housekeeping (HK) proteins. Of the 6,357 proteins identified in all 32 tissues, 1,578 proteins were classified as HK proteins (Fig. 1D); the rest showed enrichment in different tissues. Functional analysis showed the HK proteins are mainly involved in RNA processing, gene expression and protein localization, which are primary functions in all cells (Table S4).

### Protein and RNA Correlation

The correlation between RNA and protein can be defined and interpreted in different ways (*19*). Since the RNA and proteome data were generated from the same tissue specimens, we computed the correlation between protein and RNA across 32 tissues for each gene. At the RNA level, the TS-score was calculated using the same approach as for protein. The median Spearman correlation is 0.49 (interquartile range of 0.28-0.69) (Fig. 3A), consistent with previous findings (*1*–*3*). Close to half of the proteins (6,604/12,627) have statistically significant positive correlations with RNA, and, interestingly, a very small number (92) of genes showed significant negative correlations (Table S3). Among the significantly positively correlated protein/RNA, only 31% are exhibited enrichment in the same tissues. Protein and RNA which were enriched in very different tissues showed nonsignificant correlations.

**Fig. 3.**
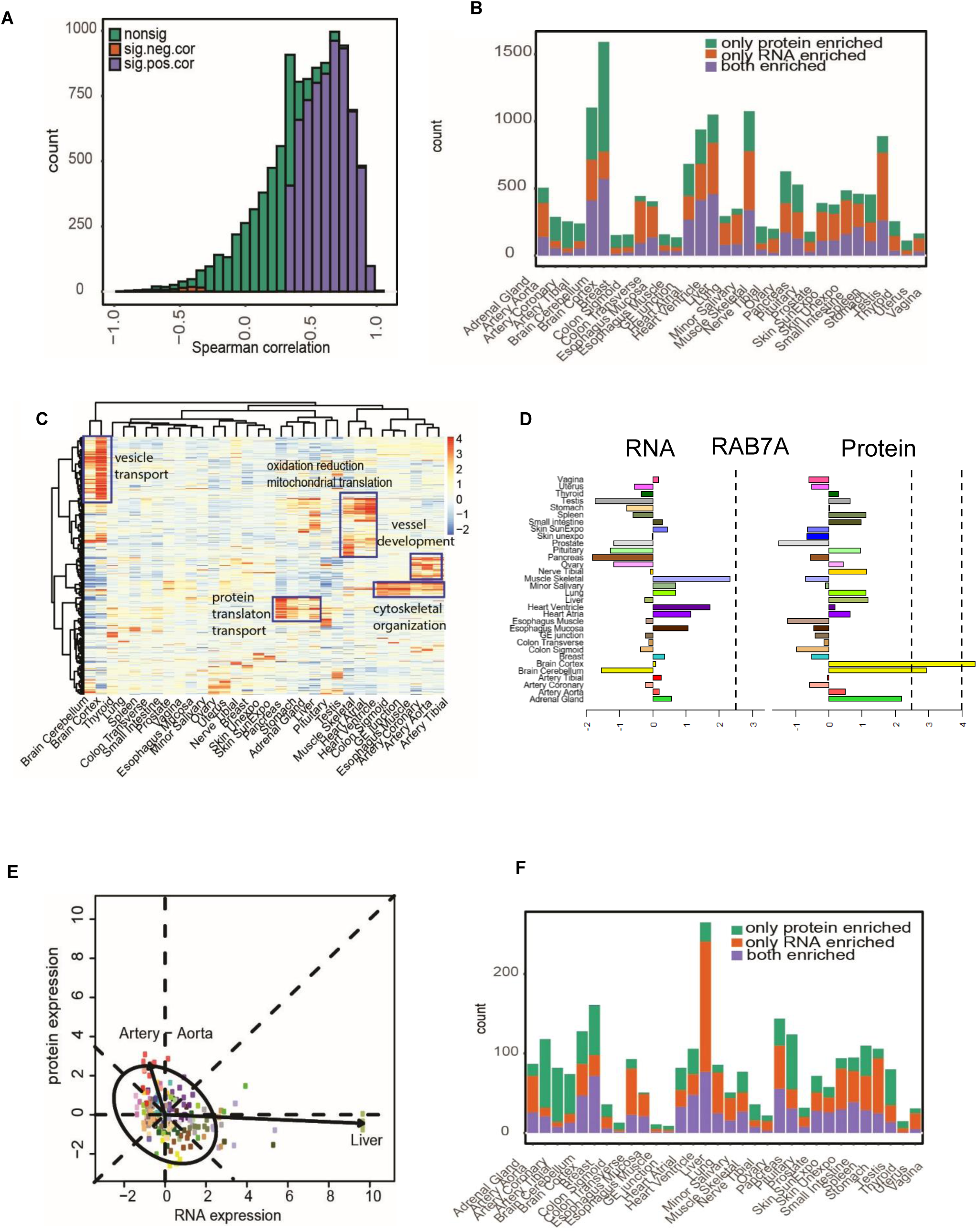
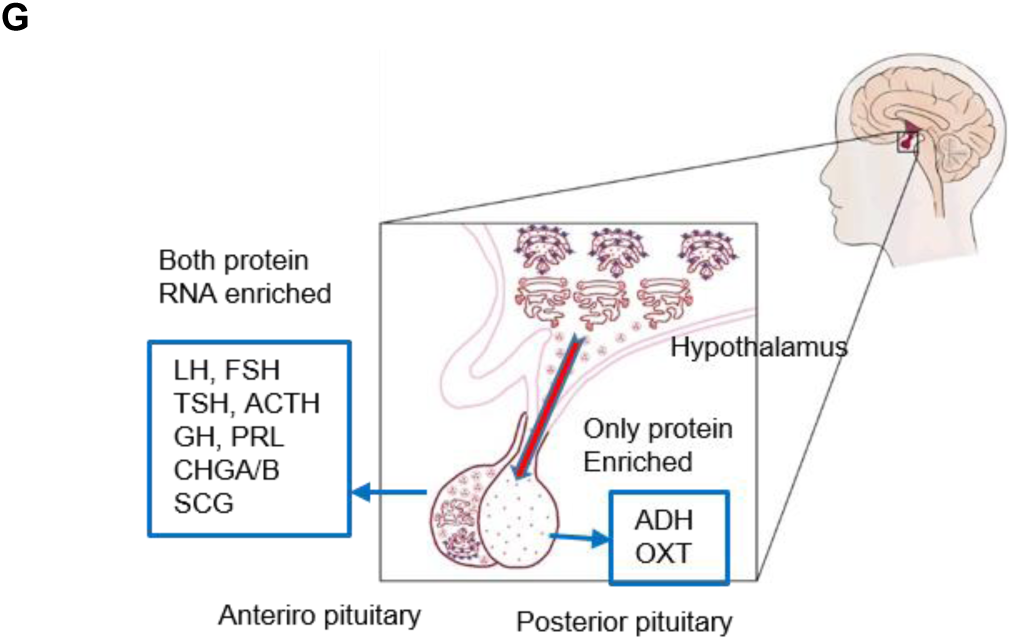
Protein and RNA correlation and concordance analysis across and within tissues. A. Spearman Correlation of protein and RNA across 32 tissues. The significance is based on permutation test from 200 permutations (Supp Sec 3.3). B. The number of concordantly and discordantly enriched proteins and RNA in each tissue. The concordance and discordance are defined in Suppl Sec 3.3. C. The enrichment of housekeeping RNAs at the protein level across tissues. D. TS-score of RAB7A across tissues in proteome and transcriptome. It is HK RNA but is enriched in brain in proteome. E. An example of two dimensional distribution of C8G at protein and RNA level across tissues. Its RNA is enriched in liver but the protein is enriched in arteries. The ellipse indicates outlier boundary (±2.5 projected into the axis) in the RNA and protein joint comparison (Supp Sec 3.3). F. Secreted proteins and their concordance to corresponding RNAs in each tissue. G. Concordance analysis of proteins secreted by pituitary. All the peptide hormones in anterior part of pituitary are concordantly enriched at protein and RNA level. Hormones in the posterior part of the pituitary are secreted from hypothalamus and stored in pituitary.

Next, we analyzed the concordance of protein and RNA enrichment in individual tissues. Concordance was defined as both protein and RNA are enriched in the same tissue as determined by outlier analysis; for discordance, only one or the other was tissue-enriched (Table S3). The concordance/discordance of every gene in each tissue can be visualized at our website (http://snyderome.stanford.edu/TSomics.html). Figure. 3B shows the number of genes concordantly and discordantly enriched in each tissue. Many proteins and RNAs are concordantly enriched; however, some are enriched only at the level of protein and not RNA. These are defined as HK genes at the RNA level. Brain has the highest number of genes only enriched at the protein level. These proteins are enriched in vesicle transport and protein localization which play important roles in neurotransmitter transport, cell-cell communication and signal transduction (Fig. 3C). Some showed extremely high enrichment with TS-score >4 such as Rab proteins which are key regulators for intracellular vesicle trafficking (Fig. 3D). In addition, many proteins in oxidation reduction pathway were also only enriched at the protein level. These proteins are essential to maintain the active function of the brain and genetic mutations affecting these proteins can cause many neurological disorders as will be discussed in disease section.

There are also many genes that are highly tissue enriched at the level of RNA but not protein. For example, testis has the highest number of genes that have RNA tissue enrichment but not protein (Table S3). Multiple proteomics studies including ours have shown that many genes expressed as RNA in testis do not have evidence of protein (*5, 20*). Failure to detect proteins may be caused by low transcription, poor translation, rapid protein degradation, limited expression or presence in only a few cell types and/or limited sensitivity of the mass spectrometer. Our data shows that at RNA TPM≤1, 1815/2756 of the RNAs are expressed in testis and 466 are only expressed in testis suggesting that lack of proteomics evidence might be caused by low RNA expression. 2,863 genes which lack detectable proteins in testis have RNA expression greater than TPM 32 where most proteins are detectable. These are likely subjected to posttranscriptional regulation accounting for their low protein abundance. Pathway analysis of the undetected proteins shows that most are involved in spermatogenesis.

Besides testis, liver also has a large number of genes that are only enriched at the RNA level. For example, for the *C8G* gene in Figure 3E, its protein abundance in liver is extremely low but the RNA level is the highest. Further analysis reveals that many of these proteins are actually secreted which will be discussed below. Pathway analysis of the rest of the discordantly RNA enriched genes showed that most are involved in mitochondria translation and cellular respiration which are not liver but heart specific functions. In fact, our data showed they are concordantly enriched in heart. Muscle also has a group of genes that are only enriched at the RNA level. These genes are involved in gene expression and mitochondrial translation. Mitochondrial ribosomal proteins (MRPSs) are enriched at the RNA level in multiple metabolic active organs such as the heart, liver and muscle but at the protein level they are only enriched in the heart. Surprisingly, other ribosomal proteins overall are discordantly enriched at the protein level, especially in pancreas, liver and stomach. Ovary is the only tissue in the opposite that only has the ribosomal RNAs enriched.

There are some genes whose RNAs are enriched in a group of tissues but their encoded proteins are selectively enriched in only a subset of them. For example, genes such as *NPPA, NPPB, MYL7* and *MYH6* are enriched at the RNA level in both heart ventricles and atria. However, at the protein level, they are enriched only in heart atria. Many genes involved in muscle contraction are enriched in both heart and skeletal muscle at the RNA level but are differentially enriched in each tissue at the protein level. The selective enrichment at the protein level indicates tissue specific functions which cannot be distinguished based on RNA information. Housekeeping genes also showed concordance/discordance at protein and RNA level. Many genes are ubiquitously expressed as HK gene at both protein and RNA level. However, some HK RNA are highly enriched in a few tissues at protein level as shown in Figure. 3C. Some HK proteins showed enrichment in a few tissues at the RNA level as well (Table S3). Overall, these results indicate differential mechanisms controlling protein and RNA levels for each gene in different tissues. How this arrangement is related to tissue function and disease is discussed below.

### Concordance and Discordance in Secreted Proteins

The protein/RNA concordance analysis indicates that some of the discordance is caused by secretion of proteins to other tissues. We systematically investigated the protein/RNA concordance in each tissue for the secreted proteins. The secreted proteins are based on HPA predictions from a combination of multiple algorithms (*4*). As described above, one would expect discordance for most secreted proteins that are constitutively secreted and target other tissues. However, our data also showed that many (501/1902) predicted secretory proteins have good protein/RNA concordance (Fig. 3F, table S3). It is likely that these proteins undergo regulated secretion in which proteins are stored in secretory vesicles and released upon stimulation (*21*). Well known examples of regulated secretion include digestive enzymes and hormones. As shown in Figure. 3F, based on the concordance and discordance results, both constitutive and regulated secretion occur in each tissue.

Among all the tissues, liver has the highest number of predicted secreted proteins followed by brain, artery, pancreas and pituitary. In liver, the largest proportion of secreted proteins are only enriched at the RNA, but not protein, level. Most of the proteins that are anti-correlated across tissues are proteins secreted from the liver. Pathway analysis shows these proteins are enriched in complement activation (*C2-9, CFHRs et.al*.), coagulation (*CFs, SERPINs*), acute phase response (*CRP, HP, ITIH4, and SAA4 et.al*), and lipid transport (apolipoproteins) and protein localization (*TF, HRG, AGT et.al*). Many of these proteins are known plasma proteins. Our data also showed the enrichment of these proteins in arteries with discordant RNA expression, which provides further evidence that these proteins are constitutively secreted to the bloodstream. However, there are some (77) liver secreted proteins that are concordantly enriched; these are mostly enzymes present in plasma that are involved in drug/amino acid metabolism and oxidation-reduction such as *CYP2* subfamily members (Table S3). Surprisingly, although secreted proteins involved in intracellular transport are not excreted, they showed high enrichment only at the protein and not RNA level. Similar enrichment discordance was observed in pancreas and brain as well. Perhaps this particular group of proteins are very stable and require only modest levels of RNA.

Similar to liver, the pancreas is a major secretory organ. Uniquely, it has both exocrine and endocrine cells that secrete many digestive enzymes and multiple hormones. Digestive enzymes in the pancreas are stored and secreted into the gut upon stimulation by food. A group of major digestive enzymes showed concordant enrichment including pancreatic amylases, lipases, proteases and others. Multiple hormones such as insulin, glucagon, chromogranins, secretogranins, progastriscin and somatostatin are also secreted by the pancreas. Insulin and glucagon are two major hormones regulated by increase/decrease of blood glucose level and are concordantly enriched at the protein and RNA level. The other hormones are discordantly lower at the protein level in the pancreas but, interestingly, are concordantly enriched in other tissues that also secrete these hormones. For example, chromogranins and secretogranins in the pituitary and progastriscin in the stomach are concordantly enriched. Unlike other hormones, these hormones are also regulated by local hormones in paracrine manner. For example, somatostatin regulates secretion of progastrin in stomach and insulin/glucagon in pancreas. This local regulation from other hormones could contribute to different concordance between RNA and protein. A few enzymes such as *SERPINAs* and *SPINT1* and other proteins such as mucin, albumin, *C4, F11* and *GC* are also enriched only at the RNA level in pancreas. Although these proteins are well known to be secreted mainly by the liver, our data suggests pancreas may also synthesize and secrete them into the bloodstream. Since the pancreas and liver share a common embryological origin and some histological similarities, it is possible they have some cellular functions in common (*22, 23*).

The pituitary is the master gland that secretes many hormones which regulate the secretion of other hormones. Our data shows hormones such as *TSH, ACTH, GH, PRL, CHGA/B, SCGs, LH* and *FSH* are all enriched in the pituitary at both the protein and RNA level. These hormones are made in the anterior part of the pituitary but are stored and undergo regulated secretion by hormones produced in the hypothalamus. Two other major hormones in the pituitary are *ADH* and *OXT*. They are both highly enriched at the protein level but not at the RNA level; these proteins are synthesized in the hypothalamus, secreted to, and stored in the posterior part of pituitary (Fig. 3G). Brain also has a group of secreted proteins that are concordantly enriched. They are not secreted into the bloodstream. Instead, most are brain specific surface proteins such as receptors for signal transduction and proteins involved in cell-cell interaction. Other tissues such as spleen and lung have a group of secreted proteins that are discordantly enriched. These proteins are mainly involved in immune response and secreted to bloodstream. In transverse colon and small intestine, proteins are secreted to lumen or extracellular matrix. The complete list of these secreted proteins with concordance analysis can be found in Table S3.

Overall, the comparison of secreted proteins provides clues as to a) which proteins undergo regulated secretion (e.g. high levels of extracellular proteins as both RNA and protein as well as b) potential sites of synthesis and action of secreted proteins (enriched only as RNA and protein, respectively).

### Metabolism

Human metabolism related diseases are emerging as global health problems. They are often complex, have multiple genes involved and different underlying mechanisms can result in similar phenotypes (*24*). Metabolic pathways are interconnected and metabolic phenotypes involve multiple organs. The exact mechanisms underlying the onset and progression of metabolism related disorders are still unclear (*25*). In the HPA study, tissue specific metabolic maps were reconstructed by integrating RNA-seq data from 32 different tissues and protein compartment information from an antibody map. Since all the metabolic reactions are carried out by enzymes, our proteomics data can provide a more direct understanding of the proteins and tissues involved in metabolism.

Proteins from 1434 genes annotated in the KEGG metabolism database were all quantified in this study. Their enrichment across 32 tissues showed that liver has the highest number of enriched metabolic proteins followed by brain, muscle and heart. To further investigate the metabolic features of each tissue, we did the enrichment test of metabolic pathways in each tissue based on the enriched proteins (Table S4). As shown in Figure. 4A, each tissue has its own unique metabolic profile. For example, in liver most (56/68) metabolic pathways are enriched except a few specific ones such as the oxidative phosphorylation pathway. This result indicates liver does not depend on the energy source from oxidative phosphorylation which is the major energy source for tissues that function aerobically. In contrast, oxidative phosphorylation, glycolysis and TCA cycle pathways are simultaneously enriched in heart, skeletal muscle and brain. Coupling of these pathways can achieve the complete oxidation of glucose and generate the maximum amount of ATP required for high energy demand. Previous studies have shown that, when active, skeletal muscle requires the most energy, whereas when resting, heart and kidney have the highest metabolism rate followed by brain and liver (*26*). Surprisingly, these pathways are also enriched in stomach which usually is not considered as a high metabolism organ. However, oxidative phosphorylation in stomach tissue is necessary for the massive acid generation by parietal cells. The stomach contains a strong acid environment that breaks down food and serves as a biological defense that eliminates pathogens, activates endothelial NADPH oxidase and increases endothelial RO (*27*). In accordance, mitochondrial proteins and enzymes involved in energy production are highly enriched in these tissues including NADH:ubiquinone oxidoreductase, ATP synthase, cytochrome c and coenzymes (Table S3). Importantly, whereas ATP synthases are enriched in muscle for energy production, the V-ATPases are specifically enriched in brain for synaptic transmission (*28*). Mutations of V-ATPase are associated with neurological diseases (*29*).

**Fig. 4.**
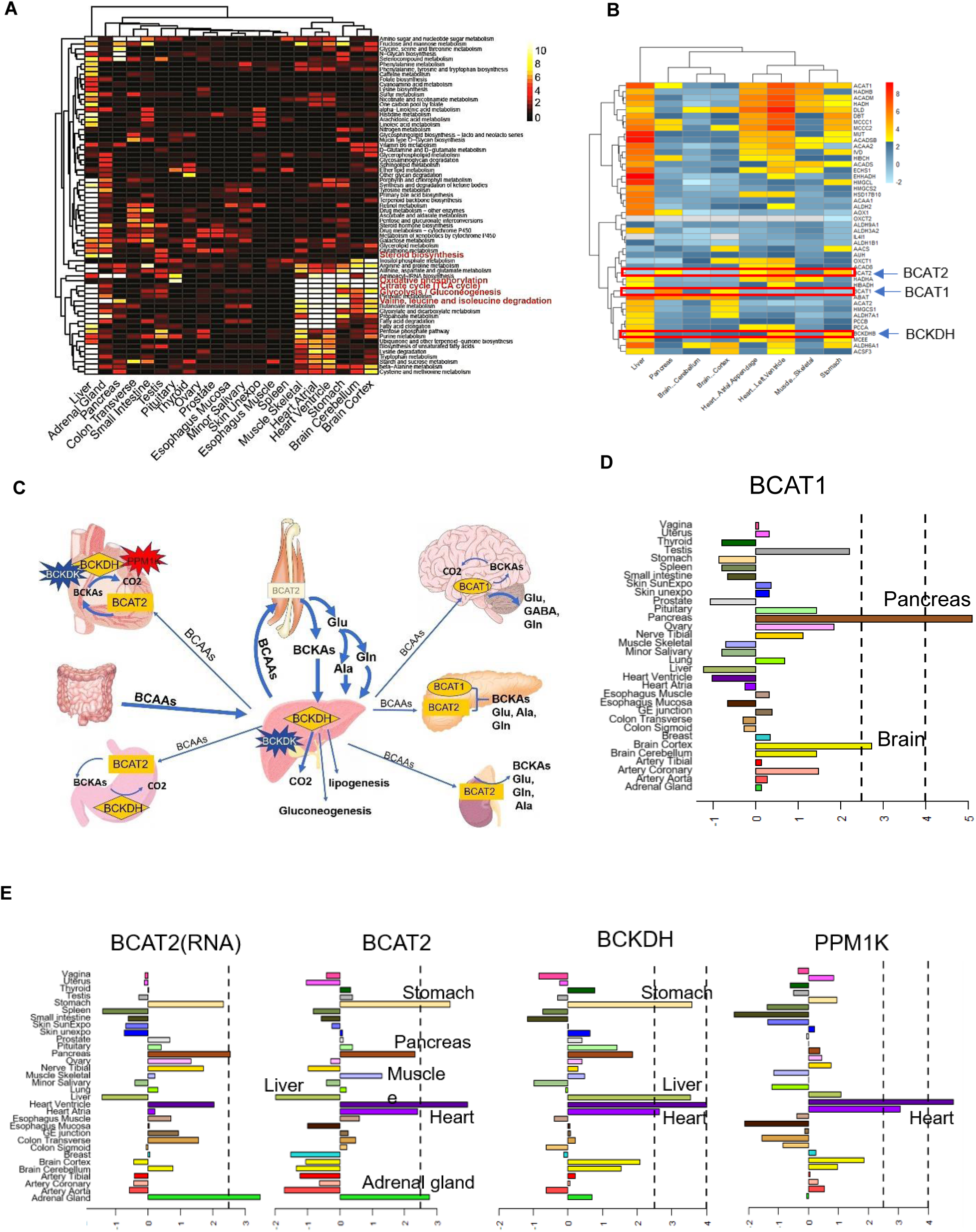
Analysis of tissue specific metabolism. A. Enrichment of metabolic pathways across different tissues. The heatmap shows the significance of the -logp values from pathway enrichment test (Suppl Sec 4 and Table S3). The plot only includes the tissues having at least one significantly enriched pathway under threshold of 0.001 for the p-value. B. The enrichment map of key enzymes in BCAA metabolism. C. Interactive map of BCAA shuttling among tissues and the enriched enzymes. D. Enrichment of BCAT1 across tissues. E. Tissue enrichment of the BCAT2 and the second step enzyme BCKDH and its activator PPM1K.

Some metabolic pathways are commonly enriched in a few tissues but they likely are compartmentalized in different parts of the cell. For example, our data shows that aminoacyl-tRNA biosynthesis pathway is enriched in heart, brain, stomach and pancreas. In heart, the high level of aminoacyl-tRNAs is mainly used for the synthesis of mitochondria proteins for high active cellular respiration, whereas in the pancreas, they are used for the active biosynthesis of proteins in the ER; thus, likely these synthetases operate in different cellular compartments. Our data showed that the steroid hormone biosynthesis pathway is not only enriched in the adrenal gland and liver but also in small intestine and transverse colon. Historically adrenal and gonad glands (ovary, testes) have been considered as major steroid hormone production organs, and liver as the main organ that metabolizes the hormones. Until recent years, evidence has shown that steroid hormones can also been produced and metabolized in other tissues such as intestine (*30*); our data are consistent with this concept.

### Branched Chain Amino Acid Metabolism

Valine, leucine and isoleucine are branched chain amino acids that play key roles in metabolism. They serve as substrates for protein synthesis or energy production and perform several metabolic and signaling functions as well. The tissue specific distribution of the BCAA metabolic enzymes, especially for the first two steps, determines the unique inter-organ BCAA metabolite shuttling (*31, 32*). In our study, most of the key enzymes involved in BCAA metabolism have been observed. Figure. 4B showed their tissue specific enrichment. Figure. 4C is a schematic view of BCAA metabolites shuttling among tissues and is annotated with the tissue enriched enzymes.

Branched-chain aminotransferase (BCAT) is the first step key metabolic enzyme with two isozymes, *BCAT1* in cytosol and *BCAT*2 in mitochondria. Our data showed that *BCAT1* is highly enriched in the pancreas followed by brain (Fig. 4D), and *BCAT2* is primarily enriched in the heart and stomach (Fig. 4E). Multiple studies have shown that the first step of BCAA metabolism mainly takes place in skeletal muscle due to the high activity of *BCAT2* (*31, 33*). *However, our data showed that BCAT2* is not enriched in skeletal muscle and its RNA level is relatively low as well (Fig. 4E). However, taking into consideration of the total weight of skeletal muscle (35-40% of body weight), moderately elevated BCAT2 level will result in substantial amount of BCAA metabolism in the initial step. Presumably, if skeleton muscle *BCAT2* level is too high, BCAAs might be mostly metabolized in skeletal muscle and cause insufficient supply to other tissues. Unlike skeletal muscle, heart has much less muscle mass and is highly enriched for *BCAT2*. Enrichment of *BCAT2* to some degree can ensure the proper amount of BCAAs needed in heart. In brain, *BCAT1* is highly enriched instead of *BCAT2*. Although these two enzymes are functionally similar, the different enrichment might suggest different roles in brain. Past studies have shown that *BCAT1* is enriched in neurons and *BCAT2* is limited to astrocytes (*34*). Since glutamate is one of the major neurotransmitters, enrichment of *BCAT1* can ensure the demand for glutamate from BCAA metabolism. The limited amount of astrocytes in bulk brain tissue might account for low protein level of *BCAT2*. In our data, pancreas is the only tissue with both forms of BCAT enriched, especially *BCAT1*. Although few studies have investigated the roles of these two enzymes in the pancreas, protein synthesis is highly active in pancreas, which could use BCAAs as substrates and energy source. Although most studies have shown that *BCAT1* is primarily expressed in brain, our data demonstrates that the *BCAT1* protein is most abundant in pancreas, in addition to its presence in the brain. Recent studies showed that BCAA level spikes years before pancreas cancer, possibly suggesting that BCAA metabolism disorder might contribute to cancer development (*35, 36*).

In the second step of BCAA metabolism, BCKAs are decarboxylated by branched-chain α-ketoacid dehydrogenase (*BCKDH*). Our data show high enrichment of *BCKDH* in heart and liver followed by stomach (Fig. 4D). The tissue enrichment information is similar to previous studies which were based on enzyme activity analysis (*37*). Among all the tissues, only heart and stomach have both *BCAT2* and *BCKDH* enriched. It has been proposed that *BCAT2* and *BCKDH* directly interact to achieve great efficiency of energy production during BCAA metabolism. The concordant enrichment of these two enzymes in heart and stomach suggests BCAA metabolism serves as an important energy source. Animal model and cardiomyocytes experiments have shown that impaired BCAA metabolism leads to the loss of cardiac contractility, premature death and induced apoptosis suggesting an important role for these enzymes in these tissues (*38, 39*).

The second step key enzyme *BCKDH* is not only regulated by its expression level in different tissues but also highly regulated by two modifying proteins, *BCKDH* kinase (*BCKDK*, inactivator) and Protein Phosphatase (*PPM1K*, activator). The imbalance of these two enzymes is involved in multiple metabolic diseases (*40*). Our data show that there is specific tissue distribution of *BCKDK* and *PPM1K* which are not always the same as *BCKDH* distribution. *BCKDK* tissue distribution is very similar to *BCKDH* distribution but the activator *PPM1K* showed extremely high enrichment in heart, but not in other tissues. A high ratio of *PPM1K* to *BCKDK* greatly favors BCKA oxidative decarboxylation. The enrichment of these enzymes likely fuels high energy production for the heart.

The downstream enzymes for further metabolization of BCAA are also highly enriched. Each intermediate metabolite during BCAA catabolism can either enter TCA cycle, used for gluconeogenesis, lipogenesis or re-aminated to form BCAA. The usage of the metabolites in each tissue depends on the homeostasis of our body and tissue specific distribution of enzymes in each step. Mutations or unbalanced amounts of enzymes may cause metabolic disorders.

### Proteome, Genetic Diseases and Drug Targets

Genetic mutations in protein coding regions can alter protein function and cause a spectrum of disease phenotypes. Since we have identified many tissue enriched proteins, we explored whether they were associated with tissue associated diseases. Figure. 5A and Table S11 shows the enrichment of disease causing proteins to a particular tissue or set of tissues. We found many cases where genetic diseases of tissue enriched proteins provided insights into the underlying disease mechanisms. For example, Bardet-Biedl syndrome (BBS) is a genetic disorder caused by mutations in at least 14 different genes which are critical to the structure and function of cilia (*41*–*43*). It is known that defects in cilia will disrupt cell movement and many chemical signaling pathways critical to many tissues. However, it is unknown exactly how each tissue is affected. BBS affiliated vision loss, polydactyly and obesity are characteristics of BBS as well as many other abnormalities such as intellectual disability, delayed motor skills and conditions that involve the heart, liver and digestive systems (*44*). Some of them can be explained by specific gene mutations but for such diverse symptoms, many are still largely unknown. In our study, proteins from 11/14 affected genes have been observed across tissues. Protein enrichment analysis shows that the majority of the BBS proteins are enriched or highest in the pituitary and some are enriched in brain, muscle or liver. Abnormality of proteins highly enriched in pituitary can cause dysfunction of pituitary which likely affect many developmental processes. One of the main symptoms such as obesity at birth could be caused by the abnormal hormone secretion. The enrichment of proteins in brain, heart, muscle and liver might contribute to related symptoms as mentioned above.

**Fig. 5.**
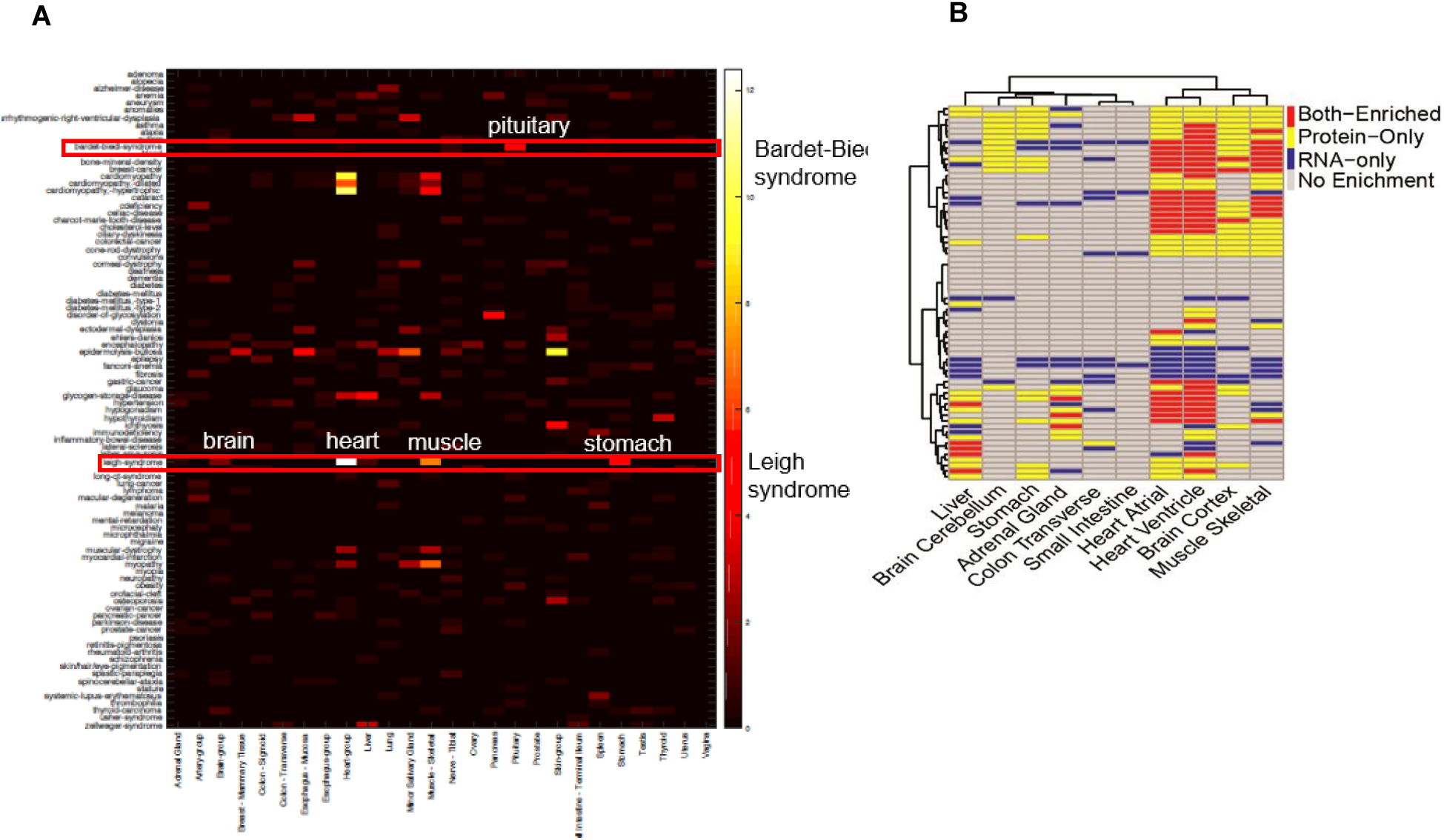
Association of tissue enriched proteins with genetic diseases. A. Heatmap of the enrichment of genetic diseases across tissues. Some genetic disease s are significantly enriched in certain tissues such as Bardet Biedl syndrome and Leigh syndrome. The disease terms are from OMIIN database. B. Protein and RNA concordance heatmap for genes involved in leigh syndrome.

Leigh syndrome is another genetic disease that is associated with mutations in as many as 75 genes. Most of the affected proteins are involved in oxidative phosphorylation in mitochondria (*45, 46*). In our study, 67/75 proteins were observed and 52 of them showed tissue enrichment. As shown in Figure. 5B, these proteins are highly enriched in a few metabolically active tissues. Heart has the highest number of enriched proteins followed by muscle, brain and stomach. Figure. 5A showed significant enrichment of the disease in these tissues. Some of these proteins were enriched in all tissues listed above and some are only enriched in one or a few. Different number of affected genes might cause different tissue related clinical symptoms. For example, the characteristic progressive loss of mental and movement abilities of this syndrome are most likely related to proteins that are enriched in brain and muscle. Some affected individuals develop hypertrophic cardiomyopathy which could be caused by mutations in proteins enriched in heart. Although stomach has the least number of enriched proteins, the first signs of Leigh syndrome seen in infancy are usually vomiting, diarrhea, and difficulty swallowing which could be explained by abnormality of stomach. Involvement of different proteins or different number of proteins might explain why a small number of individuals do not develop symptoms until adulthood or have symptoms that worsen more slowly. For the enriched proteins, we also compared their enrichment at the RNA level. As shown in Figure. 7B, protein and RNA enrichment are quite different. Although in heart and muscle some genes were concordantly enriched, almost all genes enriched at the protein level in brain and stomach did not show any enrichment at the RNA level. Thus, the specific enrichment information at protein level can help us predict the affected tissues and better understand the clinical symptoms. In this case, especially the neurological and digestive symptoms cannot be directly predicted from RNA data but could be suggested by protein enrichment information.

Many drug targets are proteins. There are 1421 identified in our data among which 466 are FDA approved drug targets and 955 are potential drug targets. 752 of the targeted proteins are enriched in certain tissues and about half are enriched in more than two tissues (Table S11). In many cases, the targeted tissue lies outside of the intended target organ suggesting possible sites of side effects. For example, valproic acid is a well known anticonvulsant exerting its effects through the inhibition of GABA transaminase (*GABAT*) in the brain as one of the main mechanisms of action. Our data showed that GABA transaminase is not only highly enriched in brain but also in the liver with the highest abundance level. The unknown liver toxicity could be caused by the inhibition of GABA transaminase in liver (*47*).

### Missing Proteins

Despite ongoing efforts to validate and annotate the proteome in normal human tissues, currently, ∼18% of proteins do not have high-stringency evidence confirming their existence. The lack of experimental data on the protein level could be caused by factors such as temporal expression, expression in tissues that are difficult to sample, low expression, or the genes do not encode functional proteins. Missing proteins are classified based on evidence of <u>protein existence</u> (PE) in a 1-5 tier system in an effort led by <u>HUPO</u> and the Human Proteome Project (*20, 48*). Since we have collected massive amounts of data from a variety of tissues, we hoped to increase the identification of missing proteins and the specific tissues where they are expressed. The identified proteins were matched to the missing protein list provided by MissingProteinPedia. In total, 310 proteins matched to the list (Table S12). Among them 87 proteins have at least 2 unique peptides (do not match to any other proteins in the current database), each with a peptide length ≥9 as required at Protein Evidence (PE) level 5. For those that do not meet these criteria, 30 have more than 2 unique peptide identifications, and 7 of them have peptide lengths longer than 20, which might add confidence to the protein identification. Surprisingly, among the 87 PE5 proteins, UNC13C and RASA4 have 11 and 17 unique peptides (≥9aa) identification, respectively. RASA4 was observed in a majority of the samples and UNC13C was identified in roughly half. UNC13C is enriched in brain and testis, and RASA4 is highly enriched in skeletal muscle. In total, 33 proteins showed enrichment in different tissues. They are enriched in brain, muscle and af few other tissues, and most are intracellular proteins. Among the 87 proteins, 33 have reliable antibody scores annotated by HPA.

### Protein Isoforms

Based on GTEx V7 RNAseq data, there are 160,907 isoforms of 22,207 genes, an average of seven isoforms per gene (*11*). The distribution of isoforms in each tissue has also been provided in the GTEx portal. Although thousands of isoforms are well documented at the transcriptome level, their expression at the protein level is still largely unknown. To identify protein isoforms, the junction peptide has to be detected by the mass spectrometer. However, due to the much lower sequence coverage compared to the RNAseq, proteome level validation of the isoform presents a substantial challenge. Previous large-scale mass spectrometry-based proteomics analyses identified only a small fraction of annotated alternative isoforms. The clearest finding from these experiments is that most human genes have a single main protein isoform (*5, 49, 50*).

In our study, proteomics data were searched against the Gencode database 19 which included 90,203 annotated protein isoforms from 15,632 annotated genes. In total, we identified 7,628 protein isoforms from 7,169 genes which have at least one unique isoform peptide (Table S5). As shown in Figure. 6A close to 7000 genes have only one protein isoform detected in our study, and very few have more than 2 isoforms, consistent with other proteomics studies (*5, 49, 50*). Since for most genes we only have one protein isoform detected, they showed similar concordance/discordance to RNA in each tissue as previous described. For proteins with more than one isoforms, some are enriched in the same tissue; for example, two isoforms of *CELA2B* are both enriched in pancreas at similar level. However, at the RNA level, one isoform is much more enriched than the other in pancreas. Some genes, such as *TPM2*, have two isoforms that are enriched in different tissues. One is enriched in heart and skeletal muscle and the other one is enriched in tissues with smooth muscles. For all the detected protein isoforms, the majority (86%) are from the most abundant RNA isoforms as shown in Figure. 6B. For the undetected protein isoforms, further analysis showed it is not mainly caused by low RNA abundance. Figure. 6C showed the proportion of protein identifications for the top two abundant RNA isoforms across different abundance levels. As shown, the detection of the rank 1 isoforms is independent of the RNA abundance level. The rank 2 isoform showed the same result except for the very low abundance ones. Comparing the proportion of protein identification between rank 1 and 2 RNA across different RNA abundance intervals showed that the detection of rank 2 protein isoform is consistently much lower than the rank 1 protein isoform (Table S9). This result suggests that the limited detection of protein isoforms is independent of their RNA abundance. The exact mechanism of selection at the protein level is still not clear and it may be caused by the limitations of current technology.

**Fig. 6.**
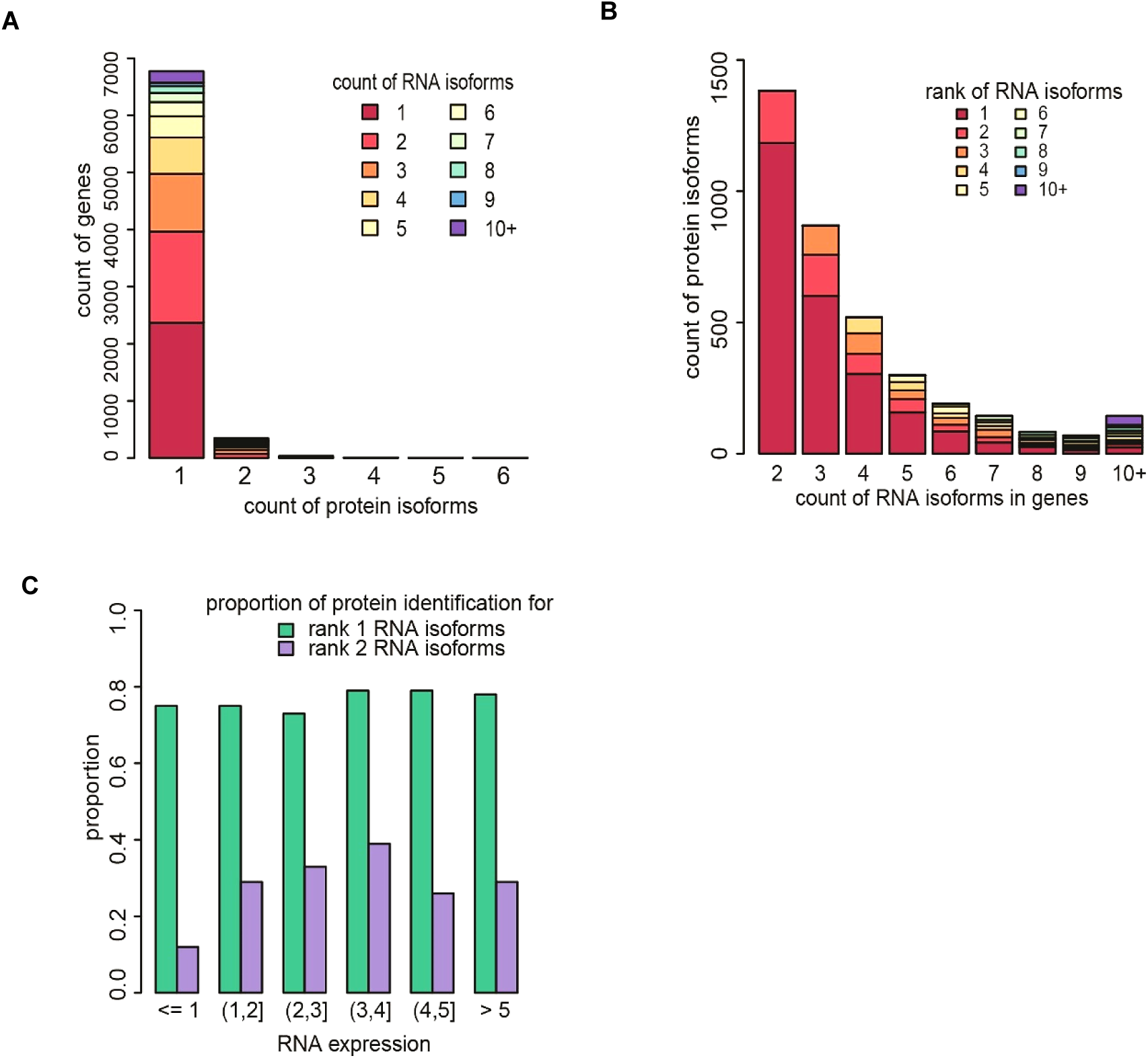
Protein isoform analysis. A. Total number of genes which have different number of isoforms identified at the protein level. Different color represents number of RNA isoforms each gene has. Although each gene has various RNA isoforms, close to 7000 gene s only has one protein isoform identified. B. The number of protein isoform identified from genes with different number of RNA isoforms. Different color represents the rank of the corresponding RNA isoform. C. The proportion of the rank 1 and 2 RNA isoforms identified at the protein level across RNA abundance intervals The chi square test is applied to test for independence of protein identification to corresponding RNA isoform expression level (Suppl Sec 4 and Table S9).

Since the majority of the genes only have one protein isoform, the diversity of tissue phenotype is probably not determined by different isoforms but differential gene expression (*50*). We investigated if there are specific enrichment of the isoforms across tissues. The enrichment analysis shows that a total of 1,826 isoforms have tissue enrichment. The RNA isoform rank of the enriched proteins in each tissue was further analyzed. We found that most of the enriched isoforms (1,655) are from the rank 1 RNA isoforms and approximately 249 enriched proteins were from the rank 2 and lower abundance isoforms (Table S5).

### Single Nucleotide Polymorphisms (SNPs)

Proteomics data obtained in our study was also used to verify the expression of SNPs at the protein level. Due to the intrinsic limitation of the mass spectrometry strategy, the chances of observing peptides harboring missense mutations are much lower than detecting variants in transcripts. However, protein information not only can provide valuable information for mutation expression but also quantitative expression of different alleles may provide insight into protein regulation. A search for peptides with missense mutations was performed using two different strategies. One matched spectra against a database with the addition of all possible missense peptides. The other used *de novo* search by PEAKS software. We intersected the two search results to filter less confident candidates. Moreover, missense peptides which were found in individuals without the corresponding genomic variant by WGS were also removed. In total, we quantitated 177 SNP peptides indicating that many SNPs are expressed. Their tissue enrichment information is listed in Table S13.

## Discussion

In this study, we have quantitatively analyzed the proteome across 32 different normal human tissues. Proteins that are highly enriched in a single tissue or a group of tissues were identified and analyzed with regards to biological function and and help provide a comprehensive map of tissue specific or shared functions. A limitation of this study is that some highly cell type-specific and/or lowly expressed proteins may be lost for identification/quantitation. However, this study is well powered to study widely expressed proteins and the variation in their abundance as well as tissue-specific proteins expressed at moderate and higher levels. Proteomics analysis also identified a group of proteins that have not been previously identified and provided evidence at the protein level for specific SNPs and isoforms. However, identification of peptides with SNPs and junction sequences of isoforms is still a significant challenge for mass spectrometry based proteomics.

The integrated proteomics and transcriptomics analysis revealed different enrichments at protein and RNA levels. Many housekeeping RNAs were enriched in multiple tissues at the protein level and a number of tissue specific RNAs, especially those expressed in testis, are not enriched in any tissue at protein level. Protein and RNA correlation across tissues showed that about half have significant positive spearman correlation. The nonsignificant correlations are primarily caused by different enrichment of protein and RNA in individual tissues. The discordance of enrichment may be due to post transcriptional or posttranslational regulation at both RNA and protein level. Our data indicated that constitutive secretion of proteins to other tissues is one major cause of the significant negative correlation and discordance of protein and RNA. The concordance analysis of all the secreted protein also stratified different secretion mechanisms (constitutive vs regulated) in each tissue. These results provide a useful resource of candidate constitutive and regulated secreted proteins and expand our understanding of regulation at omics level. All of this information can be queried and visualized on our website for a more interactive view of protein and RNA across tissues (http://snyderome.stanford.edu/TSomics.html).

The tissue specific distribution of proteins also provides an in-depth view of several complex biological events that require the interplay of multiple tissues. For example, metabolism is well coordinated by tissue specific and interconnected metabolic pathways as shown in the metabolism heatmap (Fig.4). BCAA metabolism is a unique process that requires metabolites shuttling among multiple tissues. Protein enrichment analysis provided direct evidence of the distribution of all key enzymes across tissues and shed new insights into orchestrated energy utilization. Since many complex diseases arise from the disrupted balance of metabolism, results from this multi-tissue study are expected to help us to understand the underlying mechanisms of the disease. Furthermore, for genetic diseases caused by mutations in protein coding regions, the protein enrichment information across tissues can help to predict the affected tissues and explain specific clinical symptoms. As such, the proteomic information generated in this study is expected to provide valuable insights into human biology and disease.

## Supporting information

Materials and Methods Table S1-S14 Figure S1- S16 References

## Acknowledgments

We would like to acknowledge the GTEx donors and families for donating organs to GTEx consortium for research projects. We thank Kristin Ardlie and Ellen Gelfand for coordinating the distribution and discussion of tissue samples. Thanks to Francois Aguet for constructive discussions on transcriptome data. We also Alessandra Breschi for discussion on RNAseq data analysis and Robert Tibshirani for suggestions on statistical analysis. Funding: Supported by the NIH eGTEx grant (1U01HG00761101-01) and CEGS(Center for Personal Dynamic Regulomes) grant 3P50HG00773505S1

## Author contributions

L.J. led this project in generating proteomics data, data analysis and manuscript preparation. M.W. developed statistical methods for proteomics data analysis and integration. S.L. did the initial data analysis, SNP database construction and revised the manuscript. R.J. and J.C. did proteomics sample preparation and mass spectrometry data acquisition and contributed to making figures. X.L. did the analysis on the association of tissue enriched proteins to diseases. H.F. contributed to the discussion of data analysis. G.D contributed to SNP and isoform early data analysis. H.T. and M.P.S contributed to project supervision and manuscript review and revision. All the authors contributed to manuscript revision.

## Supplementary Materials

Materials and Methods

Table S1-S14

Figure S1-S16

References

## Reference and Notes

1. N. Nagaraj, J. R. Wisniewski, T. Geiger, J. Cox, M. Kircher, J. Kelso, S. Paabo, M. Mann, Deep proteome and transcriptome mapping of a human cancer cell line. Mol. Syst. Biol. 7, 548 (2011).

2. C. Vogel, E. M. Marcotte, Insights into the regulation of protein abundance from proteomic and transcriptomic analyses. Nat. Rev. Genet. 13, 227–232 (2012).

3. S. H. Payne, The utility of protein and mRNA correlation. Trends Biochem. Sci. 40, 1–3 (2015).

4. M. Uhlen, L. Fagerberg, B. M. Hallstrom, C. Lindskog, P. Oksvold, A. Mardinoglu, A. Sivertsson, C. Kampf, E. Sjostedt, A. Asplund, I. Olsson, K. Edlund, E. Lundberg, S. Navani, C. A. Szigyarto, J. Odeberg, D. Djureinovic, J. O. Takanen, S. Hober, T. Alm, P. H. Edqvist, H. Berling, H. Tegel, J. Mulder, J. Rockberg, P. Nilsson, J. M. Schwenk, M. Hamsten, K. von Feilitzen, M. Forsberg, L. Persson, F. Johansson, M. Zwahlen, G. von Heijne, J. Nielsen, F. Ponten, Proteomics. Tissue-based map of the human proteome. Science. 347, 1260419 (2015).

5. D. Wang, B. Eraslan, T. Wieland, B. Hallström, T. Hopf, D. P. Zolg, J. Zecha, A. Asplund, L.-H. Li, C. Meng, M. Frejno, T. Schmidt, K. Schnatbaum, M. Wilhelm, F. Ponten, M. Uhlen, J. Gagneur, H. Hahne, B. Kuster, A deep proteome and transcriptome abundance atlas of 29 healthy human tissues. Mol. Syst. Biol. 15, e8503 (2019).

6. M. Wilhelm, J. Schlegl, H. Hahne, A. M. Gholami, M. Lieberenz, M. M. Savitski, E. Ziegler, L. Butzmann, S. Gessulat, H. Marx, T. Mathieson, S. Lemeer, K. Schnatbaum, U. Reimer, H. Wenschuh, M. Mollenhauer, J. Slotta-Huspenina, J. H. Boese, M. Bantscheff, A. Gerstmair, F. Faerber, B. Kuster, Mass-spectrometry-based draft of the human proteome. Nature. 509, 582–587 (2014).

7. M. S. Kim, S. M. Pinto, D. Getnet, R. S. Nirujogi, S. S. Manda, R. Chaerkady, A. K. Madugundu, D. S. Kelkar, R. Isserlin, S. Jain, J. K. Thomas, B. Muthusamy, P. Leal-Rojas, P. Kumar, N. A. Sahasrabuddhe, L. Balakrishnan, J. Advani, B. George, S. Renuse, L. D. Selvan, A. H. Patil, V. Nanjappa, A. Radhakrishnan, S. Prasad, T. Subbannayya, R. Raju, M. Kumar, S. K. Sreenivasamurthy, A. Marimuthu, G. J. Sathe, S. Chavan, K. K. Datta, Y. Subbannayya, A. Sahu, S. D. Yelamanchi, S. Jayaram, P. Rajagopalan, J. Sharma, K. R. Murthy, N. Syed, R. Goel, A. A. Khan, S. Ahmad, G. Dey, K. Mudgal, A. Chatterjee, T. C. Huang, J. Zhong, X. Wu, P. G. Shaw, D. Freed, M. S. Zahari, K. K. Mukherjee, S. Shankar, A. Mahadevan, H. Lam, C. J. Mitchell, S. K. Shankar, P. Satishchandra, J. T. Schroeder, R. Sirdeshmukh, A. Maitra, S. D. Leach, C. G. Drake, M. K. Halushka, T. S. Prasad, R. H. Hruban, C. L. Kerr, G. D. Bader, C. A. Iacobuzio-Donahue, H. Gowda, A. Pandey, A draft map of the human proteome. Nature. 509, 575–581 (2014).

8. M. Beck, A. Schmidt, J. Malmstroem, M. Claassen, A. Ori, A. Szymborska, F. Herzog, O. Rinner, J. Ellenberg, R. Aebersold, The quantitative proteome of a human cell line. Mol. Syst. Biol. 7, 549 (2011).

9. G. Kempermann, Faculty of 1000 evaluation for The Genotype-Tissue Expression (GTEx) pilot analysis: Multitissue gene regulation in humans. F1000 - Post-publication peer review of the biomedical literature (2015),, doi: 10.3410/f.725479865.793506594.

10. E. Project, eGTEx Project, Enhancing GTEx by bridging the gaps between genotype, gene expression, and disease. Nature Genetics. 49 (2017), pp. 1664– 1670.

11. M. Mele, P. G. Ferreira, F. Reverter, D. S. DeLuca, J. Monlong, M. Sammeth, T. R. Young, J. M. Goldmann, D. D. Pervouchine, T. J. Sullivan, R. Johnson, A. V. Segre, S. Djebali, A. Niarchou, T. G. Consortium, F. A. Wright, T. Lappalainen, M. Calvo, G. Getz, E. T. Dermitzakis, K. G. Ardlie, R. Guigo, The human transcriptome across tissues and individuals. Science. 348 (2015), pp. 660–665.

12. M. Barnes, Faculty of 1000 evaluation for Genetic effects on gene expression across human tissues. F1000 - Post-publication peer review of the biomedical literature (2017),, doi: 10.3410/f.731947328.793537670.

13. L. J. Carithers, H. M. Moore, The Genotype-Tissue Expression (GTEx) Project. Biopreservation and Biobanking. 13 (2015), pp. 307–308.

14. G. C. McAlister, D. P. Nusinow, M. P. Jedrychowski, M. Wühr, E. L. Huttlin, B. K. Erickson, R. Rad, W. Haas, S. P. Gygi, MultiNotch MS3 enables accurate, sensitive, and multiplexed detection of differential expression across cancer cell line proteomes. Anal. Chem. 86, 7150–7158 (2014).

15. T. Geiger, A. Velic, B. Macek, E. Lundberg, C. Kampf, N. Nagaraj, M. Uhlen, J. Cox, M. Mann, Initial quantitative proteomic map of 28 mouse tissues using the SILAC mouse. Mol. Cell. Proteomics. 12, 1709–1722 (2013).

16. L. Wu, S. I. Candille, Y. Choi, D. Xie, L. Jiang, J. Li-Pook-Than, H. Tang, M. Snyder, Variation and genetic control of protein abundance in humans. Nature. 499, 79–82 (2013).

17. M. Wang, L. Jiang, R. Jian, J. Chen, M. P. Snyder, H. Tang, RobNorm: Model-Based Robust Normalization for High-Throughput Proteomics from Mass Spectrometry Platform. bioRxiv (2019), p. 770115.

18. D. Hartl, R. Tirouvanziam, J. Laval, C. M. Greene, D. Habiel, L. Sharma, A. Ö. Yildirim, C. S. Dela Cruz, C. M. Hogaboam, Innate Immunity of the Lung: From Basic Mechanisms to Translational Medicine. J. Innate Immun. 10, 487–501 (2018).

19. Y. Liu, A. Beyer, R. Aebersold, On the Dependency of Cellular Protein Levels on mRNA Abundance. Cell. 165, 535–550 (2016).

20. F. Jumeau, E. Com, L. Lane, P. Duek, M. Lagarrigue, R. Lavigne, L. Guillot, K. Rondel, A. Gateau, N. Melaine, B. Guével, N. Sergeant, V. Mitchell, C. Pineau, Human Spermatozoa as a Model for Detecting Missing Proteins in the Context of the Chromosome-Centric Human Proteome Project. Journal of Proteome Research. 14 (2015), pp. 3606–3620.

21. A. Feizi, F. Gatto, M. Uhlen, J. Nielsen, Human protein secretory pathway genes are expressed in a tissue-specific pattern to match processing demands of the secretome. NPJ Syst Biol App l. 3, 22 (2017).

22. E. Ghurburrun, I. Borbath, F. P. Lemaigre, P. Jacquemin, Liver and Pancreas: Do Similar Embryonic Development and Tissue Organization Lead to Similar Mechanisms of Tumorigenesis? Gene Expr. 18, 149–155 (2018).

23. M. Esrefoglu, E. Taslidere, A. Cetin, Development of Liver and Pancreas. Bezmialem Science. 5 (2016), pp. 30–35.

24. J. J. Heindel, B. Blumberg, M. Cave, R. Machtinger, A. Mantovani, M. A. Mendez, A. Nadal, P. Palanza, G. Panzica, R. Sargis, L. N. Vandenberg, F. Vom Saal, Metabolism disrupting chemicals and metabolic disorders. Reprod. Toxicol. 68, 3– 33 (2017).

25. C. Angione, Human Systems Biology and Metabolic Modelling: A Review—From Disease Metabolism to Precision Medicine. Biomed Res. Int. 2019 (2019), doi: 10.1155/2019/8304260.

26. Z. Wang, Z. Ying, A. Bosy-Westphal, J. Zhang, B. Schautz, W. Later, S. B. Heymsfield, M. J. Müller, Specific metabolic rates of major organs and tissues across adulthood: evaluation by mechanistic model of resting energy expenditure. Am. J. Clin. Nutr. 92, 1369–1377 (2010).

27. H. Suzuki, T. Nishizawa, H. Tsugawa, S. Mogami, T. Hibi, Roles of oxidative stress in stomach disorders. J. Clin. Biochem. Nutr. 50, 35–39 (2012).

28. A. Bodzeta, M. Kahms, J. Klingauf, The Presynaptic v-ATPase Reversibly Disassembles and Thereby Modulates Exocytosis but Is Not Part of the Fusion Machinery. Cell Reports. 20 (2017), pp. 1348–1359.

29. A. Fassio, A. Esposito, M. Kato, H. Saitsu, D. Mei, C. Marini, V. Conti, M. Nakashima, N. Okamoto, A. Olmez Turker, B. Albuz, C. N. Semerci Gündüz, K. Yanagihara, E. Belmonte, L. Maragliano, K. Ramsey, C. Balak, A. Siniard, V. Narayanan, C4RCD Research Group, C. Ohba, M. Shiina, K. Ogata, N. Matsumoto, F. Benfenati, R. Guerrini, De novo mutations of the ATP6V1A gene cause developmental encephalopathy with epilepsy. Brain. 141, 1703–1718 (2018).

30. H. Wittenburg, U. Tennert, J. Mössner, Hormonal and metabolic functions of the small intestine. Internist. 51, 695–701 (2010).

31. M. Holeček, Branched-chain amino acids in health and disease: metabolism, alterations in blood plasma, and as supplements. Nutr. Metab.. 15, 33 (2018).

32. Z. Arany, M. Neinast, Branched Chain Amino Acids in Metabolic Disease. Curr. Diab. Rep. 18, 76 (2018).

33. S. M. Hutson, A. J. Sweatt, K. F. LaNoue, Branched-Chain Amino Acid Metabolism: Implications for Establishing Safe Intakes. The Journal of Nutrition. 135 (2005), p. 1557S–1564S.

34. J. E. Sperringer, A. Addington, S. M. Hutson, Branched-Chain Amino Acids and Brain Metabolism. Neurochem. Res. 42, 1697–1709 (2017).

35. R. Katagiri, A. Goto, T. Nakagawa, S. Nishiumi, T. Kobayashi, A. Hidaka, S. Budhathoki, T. Yamaji, N. Sawada, T. Shimazu, M. Inoue, M. Iwasaki, M. Yoshida, Tsugane, Increased Levels of Branched-Chain Amino Acid Associated With Increased Risk of Pancreatic Cancer in a Prospective Case–Control Study of a Large Cohort. Gastroenterology. 155, 1474–1482.e1 (2018).

36. J. R. Mayers, C. Wu, C. B. Clish, P. Kraft, M. E. Torrence, B. P. Fiske, C. Yuan, Y. Bao, M. K. Townsend, S. S. Tworoger, S. M. Davidson, T. Papagiannakopoulos, A. Yang, T. L. Dayton, S. Ogino, M. J. Stampfer, E. L. Giovannucci, Z. R. Qian, D. A. Rubinson, J. Ma, H. D. Sesso, J. M. Gaziano, B. B. Cochrane, S. Liu, J. Wactawski-Wende, J. E. Manson, M. N. Pollak, A. C. Kimmelman, A. Souza, K. Pierce, T. J. Wang, R. E. Gerszten, C. S. Fuchs, M. G. Vander Heiden, B. M. Wolpin, Elevation of circulating branched-chain amino acids is an early event in human pancreatic adenocarcinoma development. Nat. Med. 20, 1193–1198 (2014).

37. S. E. Gillim, R. Paxton, G. A. Cook, R. A. Harris, Activity state of the branched chain α-ketoacid dehydrogenase complex in heart, liver, and kidney of normal, fasted, diabetic, and protein-starved rats. Biochem. Biophys. Res. Commun. 111, 74–81 (1983).

38. Y. Huang, M. Zhou, H. Sun, Y. Wang, Branched-chain amino acid metabolism in heart disease: an epiphenomenon or a real culprit? Cardiovasc. Res. 90, 220–223 (2011).

39. X. Du, Y. Li, Y. Wang, H. You, P. Hui, Y. Zheng, J. Du, Increased branched-chain amino acid levels are associated with long-term adverse cardiovascular events in patients with STEMI and acute heart failure. Life Sci. 209, 167–172 (2018).

40. P. J. White, R. W. McGarrah, P. A. Grimsrud, S.-C. Tso, W.-H. Yang, J. M. Haldeman, T. Grenier-Larouche, J. An, A. L. Lapworth, I. Astapova, S. A. Hannou, George, M. Arlotto, L. B. Olson, M. Lai, G.-F. Zhang, O. Ilkayeva, M. A. Herman, R. M. Wynn, D. T. Chuang, C. B. Newgard, The BCKDH Kinase and Phosphatase Integrate BCAA and Lipid Metabolism via Regulation of ATP-Citrate Lyase. Cell Metab. 27, 1281–1293.e7 (2018).

41. N. Haq, C. Schmidt-Hieber, F. J. Sialana, L. Ciani, J. P. Heller, M. Stewart, L. Bentley, S. Wells, R. J. Rodenburg, P. M. Nolan, E. Forsythe, M. C. Wu, G. Lubec, P. Salinas, M. Häusser, P. L. Beales, S. Christou-Savina, Loss of Bardet-Biedl syndrome proteins causes synaptic aberrations in principal neurons. PLoS Biol. 17, e3000414 (2019).

42. S. A. Khan, N. Muhammad, M. A. Khan, A. Kamal, Z. U. Rehman, S. Khan, Genetics of human Bardet-Biedl syndrome, an updates. Clinical Genetics. 90 (2016), pp. 3–15.

43. K. E. Knockenhauer, T. U. Schwartz, Bardet-Biedl Syndrome 9 Protein (aa1-407), Homo sapiens (2015),, doi: 10.2210/pdb4yd8/pdb.

44. L. Foggensteiner, P. Beales, Bardet–Biedl syndrome and other ciliopathies. Oxford Medicine Online (2015),, doi: 10.1093/med/9780199592548.003.0314.

45. S. Dimauro, D. C. de Vivo, Genetic heterogeneity in leigh syndrome. Ann. Neurol. 40, 5–7 (1996).

46. N. J. Lake, A. G. Compton, S. Rahman, D. R. Thorburn, Leigh syndrome: One disorder, more than 75 monogenic causes. Ann. Neurol. 79, 190–203 (2016).

47. V. Gayam, A. K. Mandal, M. Khalid, B. Shrestha, P. Garlapati, M. Khalid, Valproic acid induced acute liver injury resulting in hepatic encephalopathy-a case report and literature review. Journal of Community Hospital Internal Medicine Perspectives. 8 (2018), pp. 311–314.

48. M. S. Baker, S. B. Ahn, A. Mohamedali, M. T. Islam, D. Cantor, P. D. Verhaert, S. Fanayan, S. Sharma, E. C. Nice, M. Connor, S. Ranganathan, Accelerating the search for the missing proteins in the human proteome. Nat. Commun. 8, 14271 (2017).

49. I. Ezkurdia, J. M. Rodriguez, E. Carrillo-de Santa Pau, J. Vázquez, A. Valencia, M. L. Tress, Most highly expressed protein-coding genes have a single dominant isoform. J. Proteome Res. 14, 1880–1887 (2015).

50. M. L. Tress, F. Abascal, A. Valencia, Alternative Splicing May Not Be the Key to Proteome Complexity. Trends Biochem. Sci. 42, 98–110 (2017).

