## Supplementary material for "A Quantitative Proteome Map of the Human Body": Materials and Methods Table S1-S14 Figure S1- S16 References

**This file includes:**

Materials and Methods

Figs. S1 to S16

Tables S6 to S9 and S14

Captions for data tables S1 to S5 and S10 to S13

References

**Supplementary Material for**

**“A Quantitative Proteome Map of the Human Body”**

Lihua Jiang, Meng Wang, Shin Lin, Ruiqi Jian, Xiao Li, Joanne Y. Chan, Huaying Fang,

GTEx Consortium, Hua Tang*, Michael P. Snyder*.

**Table of Contents**

1. **Sample preparation and mass spectrometry workflow**

1.1 Sample preparation

1.2 Mass Spectrometry Data Acquisition and Analysis

2. **Data processing**

2.1 Protein quantification across tissues

2.2 RNA quantification across tissues

2.3 Isoform quantification in protein and RNA

3. **Tissue specificity score**

3.1 Tissue specificity score in protein expression

3.2 Tissue specificity score in RNA expression

3.3 Tissue specificity score comparison in protein and RNA

3.4 Tissue specificity score in isoforms

**4. Methods in the analysis**

**Materials and Methods**

**1.** **Sample collection and experimental design**

**1.1 Sample Preparation**

Paxgene fixed tissue samples were provided by NIH GTEx consortium. Detailed information about donor enrollment, tissue collection, sample fixation and  histopathological review methods are described in (*1, 2*). There are in total 201 samples from 32 major organs from 14 different individuals. The sample preparation method was as described before with modification *(3).* About 30mg tissue samples were cut into small pieces on ice and further disrupted using beat beating and sonication in lysis buffer (6 M guanidine, 10mM TCEP, 40mM CAA, 100mM Tris pH 8.5). The supernatant was collected and heated at 95$℃$ for 5min. After protein reduction and alkylated, protein concentration was measured using the BCA kit (ThermoFisher). Since Paxgene fixed samples have a high amount of PEG contamination, protein extract was cleaned up by acetone precipitation at -20⁰C overnight. The protein pellet was washed with acetone 3 times and air-dried. The pellet was resuspended in 6 M guanidine and 100mg was used for digested using LysC (1:100 protease to protein ratio) for 2 hours followed by trypsin (1:50) digestion overnight at 37⁰C. Peptides were cleaned up using Waters HLB column and subsequently labeled using TMT10 Plex (ThermoFisher) in 100mM TEAB buffer. An equal amount of proteins from each tissue were pooled together as a reference sample. Tissue samples were randomized and equal amount of them and one common reference sample was multiplexed into one sample. To ensure equal mix, we mixed a small amount of each sample first and adjusted the amount of each sample for the final run based on the mass spectrometry results of the small mix.

About 15ug of multiplexed sample was loaded to Waters 2D LC system for online fractionation. Peptides were separated by reverse-phase chromatography at high pH in the first dimension, followed by an orthogonal separation at low pH in the second dimension. In the first dimension, the mobile phases were buffer A: 20mM ammonium formate at pH10 and buffer B: Acetonitrile. Peptides were separated on an Xbridge 300µm x 5 cm C18 5.0µm column (Waters) using 12 discontinuous step gradient at 2 µl/min. In the second dimension, peptides were loaded to an in-house packed 75µm ID/15µm tip ID x 25cm Sepax GP-C18 1.8µm resin column with buffer A (0.1% formic acid in water). Peptides were separated with a linear gradient from 5% to 30% buffer B (0.1% formic acid in acetonitrile) at a flow rate of 300 nl/min in 180 min. The LC system was directly coupled in-line with an Orbitrap Fusion (Thermo Fisher Scientific).

**1.2 Mass Spectrometry Data Acquisition and Analysis**

The Orbitrap Fusion was operated in a data-dependent mode for both MS2 and MS3. MS1 scan was acquired in the Orbitrap mass analyzer with resolution 120,000 at m/z 400. Top speed instrument method was used for MS2 and MS3. For MS2, the isolation width was set at 0.7 Da and isolated precursors were fragmented by CID at a normalized collision energy (NCE) of 35% and analyzed in the ion trap using “turbo” scan. Following the acquisition of each MS2 spectrum, a synchronous precursor selection (SPS) MS3 scan was collected on the top 5 most intense ions in the MS2 spectrum. SPS-MS3 precursors were fragmented by higher energy collision-induced dissociation (HCD) at an NCE of 65% and analyzed using the Orbitrap at a resolution of 60,000.

We used SEQUEST in ProteomeDiscoverer (ThermoFisher Scientific) for protein identification. Raw files from 12 fractions of each sample were combined together for a single search against GENCODE V19 human proteome database (*4*). Mass tolerance of 10ppm was used for precursor ion and 0.6 Dalton for fragment ions. The search included cysteine carbamidomethylation as a fixed modification. Peptide N-terminal and lysine TMT 10plex modification, protein N-terminal acetylation and methionine oxidation were set as variable modifications. Up to two missed cleavages were allowed for trypsin digestion. The peptide false discovery rate (FDR) was set as <1% using Percolator. For protein identification, at least one unique peptide with a minimum 6 amino acid length was required. For protein quantitation, only unique peptides with reporter ion mass tolerance of less than 10ppm were used. Peptide precursor ion isolation purity should be >50%, signal-to-noise (S/N) > 15 and the summed S/N of all channels > 200. Peptides passing these criteria were summed, thereby giving more weight to the most-intense peptides. We also pooled together all the spectra in this study for a single search at protein FDR of 1%. For structure variant peptide search, we reconstructed the protein database by adding all the structure variant peptides to the database. The structure variant peptides were extracted based on the SNP information provided by GTEx consortium.

**2.** **Data processing**

**2.1 Protein quantification across tissues**

**Quantification of each gene at the protein level**

Protein abundance of each sample was first rescaled so that the total sum of the peptide abundance in each channel was the same as the average of the total sum of abundance of the two reference channels in the same run. In each 10plex sample, if a peptide abundance in one channel is less than 15 and the total sum of 10 channels is less than 200, its abundance was set as NA. The abundance of peptides that are unique to a gene was summed to represent the protein abundance level of the gene. In total, we identified proteins from 13,813 genes and quantified 12,627 of them after peptide level filtering. The protein quantitative information combined from 56 runs is shown in Table S1.

**Robust normalization**

Since we have a common reference sample in each run, batch effects were removed by using the relative abundance of each sample to the reference sample. NAs in the reference channels (126, 131 channels) were imputed using a minimum value of 15. The relative abundance of each sample was logarithm transformed at base 2. Different from traditional case-control study or a study with a few conditions, our samples are from 32 different types of tissues, which are highly heterogeneous. Majority of previous normalization methods cannot guarantee a robust and tissue adaptive correction. Here, we used a data-driven robust normalization method (RobNorm) which weighted in tissue sample heterogeneities (*5*). To robustly estimate the sample effects, we implemented the density power weight to down weigh the outliers for our structured data. Our algorithm automatically detects the sample inliers which was subsequently used for the robust normalization and at the same time keeps the heterogeneities in outliers. To avoid the bias from missing values, the estimation of sample effects was based on the genes with less than 50% missing values, in total, 6,320 genes. We set density power parameter $\gamma=1$, and took zero vector as the standard sample in RobNorm (*5*). The sample effects were corrected on the relative abundances of all genes from 420 samples (two technical replicates). After normalization, the log ratio values were transformed back to the absolute abundances for missing value imputation in the next step. The box plots of relative abundances in log scale of tissue samples before and after robust normalization are shown in Fig.S1 (a-b). The normalized absolute abundances in the protein profile are summarized in Table S1.

**Missing value imputation**

Within-run missing value due to low expression was imputed with the minimum expression of 15. If a protein was missing across different runs, no value was imputed. However, for highly tissue specific proteins, they will not be detected in multiple runs without the specific tissue in the sample. In this case, we applied a significance test to detect if the highly missing proteins (occurred in < 28 runs) are associated with a specific tissue. The protein was considered to have a run occurrence if in at least one tissue its abundance > 15. Fisher’s exact test was applied for testing whether the run occurrences of a protein are enriched in a certain tissue, whether its occurrences are tissue-driven. The p-values was adjusted by Bonferroni’s correction. If a protein has minimum p-values (from 32 tissues) < 0.2/32, we call this protein tissue-driven occurrence protein (871 proteins) and the tissue as a driven tissue. For these proteins, the missing value was first imputed by the minimum abundance. According to the following steps, we then calculated tissue specificity (TS) scores for these highly missing proteins. If at least one driven tissue has TS score $\geq$ 2.5, this protein is not only tissue-driven but also tissue enriched/specific and thus the across-run imputation is implemented. If none of the “driven” tissue(s) has TS score $\geq$ 2.5, this protein will not get across-run imputation to proceed the rest steps (169 proteins).

**Tissue level filtering and technical replicates combination**

After robust normalization and tissue sample imputation, we then took the non-missing absolute expression < 15 to be 15 and obtained the relative abundance in log scale at base 2. To combine technical replicates, we first did tissue level filtering. For each gene and each tissue, if a technical replicate is 2.5 median absolute deviation (MAD) away from the corresponding tissue median, its value was set as NA. To maintain the individual variation, if all replicates from the same person are outliers, we still kept those replicates. After filtering, we took an average of technical replicates for each gene and each tissue and finally constructed a protein expression matrix from 12,628 genes and 201 samples. The following tissue specificity analysis was based on this expression matrix, which is summarized in Table S1. The hierarchical plot of protein tissue samples is shown in Fig.S2.

**Tissue level abundance**

Based on the technical combined filtered dataset, we took the sample median on the protein expressions (in log scale) for each tissue as tissue level abundance for each gene, which is summarized in Table S1.

**2.2 RNA quantification across tissues**

**Dataset**

We obtained the RNAseq data from [GTEx portal](https://gtexportal.org/) in version 7 based on the GENCODE v19 annotation. GTEx v7 data includes 18,777 protein-coding genes from 11,688 samples, where 12,238 genes are quantified at the protein level in our study. We analyzed 182 RNA-seq data corresponding to the samples from 14 individuals in our study.

There are 1,155 genes with the tissue median RNA TPM less than 1 based on the large cohort data. They are excluded from the tissue enrichment analysis. Among all the protein-coding genes, there are 9,413 genes present in all tissues with median TPMs > 1 based on the large cohort data. Out of 9,413 genes, 7,855 are quantified at both RNA and protein level.

**Robust normalization**

To avoid taking the log of zero TPM (Transcripts Per Kilobase Million), a small value was added to genes with TPM close to zero. When a TPM lies within the range of $[0, 0.01]$, it was replaced with a value uniformly from $[0.001, 0.01]$. Next, all the values were log transformed at base 2. For 14 subjects in our study, there are 8,083 genes with raw tissue median TPMs > 1. Based on those genes, we applied our normalization method RobNorm in (*5*) on the log of TPM. The sample effect was estimated under density power parameter $\gamma=1$ and the sample medians were used as the standard sample to adjust for the RNA profile of all the protein-coding genes. The box plots of RNA abundances of tissue samples before and after robust normalization are shown in Fig.S3 (a-b). The hierarchical plot of RNA tissue samples is shown in Fig.S4.

**Tissue level abundance**

We took sample median on the normalized RNA expressions (in log scale) for each tissue as tissue level abundance for each gene, which is summarized in Table S2.

**2.3 Isoform quantification in protein and RNA**

Each isoform has to have at least one unique peptide to the isoform. In total, we identified 7,628 protein-coding isoforms corresponding to 7,169 genes. To quantify isoforms, using the same filter criteria on the peptides as in the gene level, we quantified 6,876 protein isoforms corresponding to 6,538 genes. We applied the same procedures as in gene level in Section 2.1 to obtain isoform protein expressions.

In the RNA level, due to the fact that some isoforms sharing the same CDs are called as different isoforms, we collapsed the transcripts IDs with the same CDs based on Gencode v19. The identified protein-coding isoform names in both protein and RNA are summarized in Table S4. We retrieved isoform expression from [GTEx portal](https://gtexportal.org/) in version 7 and applied the same procedures as in gene level in Section 2.2 to obtain isoform RNA expressions.

**3. Tissue specificity score**

**3.1 Tissue specificity score in protein expression**

**Protein population**

In previous studies, several methods have been developed to define tissue specificity (TS) scores *(6)*. As discussed in *(6)*, the key for defining tissue specificity is to distinguish inlier and outlier tissues. Due to the complexity of the tissue-specific outliers, we focused on the inliers and defined the concept of expression population. When comparing samples from multiple tissues in our data, the main effect in the population is the tissue effect, which is confirmed by the t-SNE plot *(7)* in Fig.4A in the main text and hierarchical cluster plot in Fig.S2, that is, the tissue samples are clustered by tissue types. Fig.S5 showed the distribution of the protein abundance across tissues for 16 randomly selected proteins without missing values. For each protein, its abundance distribution across tissues appeared as a main density bump (population distribution) with some specific tissues outside the population. In statistics, it is a challenge to determine population distribution especially under the presence of outliers *(8-10)* and in biology, it is more challenging when there is no ground truth. Based on our data and results from several other works *(11, 12)*, we considered the population distribution as Gaussian. Noting the gene heterogeneities, we developed a novel data-adaptive and robust estimation method for the population fitting, where the data itself can select a tuning parameter to adapt its heterogeneous expressions and thus better fit the population. The statistical analysis for this procedure can be found in AdaReg *(13)*. Its application and comparison to other methods are discussed in AdaTiSS *(6)*.

We applied our population fitting algorithm (AdaTiSS) on the preprocessed data to obtain population mean, population standard deviation and population proportion. Due to the presence of missing values, we only fit the population for the genes that were observed (after imputation) in at least 50 samples (out of 201) and at least 50 samples away from zero (since from our imputation step, there could be imputed many zeros), in total 8,412 genes (out of 12,627 quantified proteins). For the rest of the proteins, we obtained population mean from sample median, population standard deviation from sample MAD, and population proportion from the proportion of the observed samples within 2MAD from the median. As the cases of the proteins having zero MAD, we took the population standard deviation of less than 0.01 to be 0.01. Fig.S6 shows the histogram of the fitted population in protein from our algorithm. All the population fitting parameters in protein can be found in Table S3.

As a comparison to the standard sample median and sample MAD, Fig.S7 shows gene-wise comparisons for the population mean and population standard deviation (SD) on the 8,412 genes fitted using our algorithm. There are 85 genes whose ratio of our fitted SD over MAD is greater than 1.2 and 777 genes whose ratio is less than 0.8. Fig.S8 shows histograms of protein expression for 16 randomly selected genes whose SD ratio is between 0.8 and 1.2, where the fitted density is a nice fit for the big density bump. Fig.S9 shows histograms for the genes whose SD ratio is less than 0.8, i.e., the MAD is larger. We can see there are a bunch of samples lying in the tails of the big bump. We think this is one advantage of our method maintaining robustness under heavy outliers. It could happen there are two close density bumps shown in some genes. In this case, our algorithm tends to fit one single density. Fig.S10 shows histograms for the genes whose SD ratio is greater than 1.2, i.e., our fitted SD is larger. Such cases mainly occur when there is a local concentration that may come from our imputation step of imputing some low expressions by 15 leading the population fitting challenge. More discussions and details to see in AdaReg *(13)*.and AdaTiSS *(6)*. As a resource, we provide all the population information and population fitting on our website.

**Protein TS scores**

Once we robustly captured the population, the TS is calculated based on the robust z-score. Suppose $X$ is the expression matrix with genes in rows and samples in columns. Define the sample robust z-score $Z$ for gene $i$ sample $j$ by

$$Z_{ij}=\frac{X_{ij}-{(population mean)}_{i}}{{(population standard deviation)}_{i}},$$

and the TS score tissue $t$ by

$$S_{it}= {median}_{j_{t}}Z_{ij},$$

where $j_{t}$ is the sample $j$’s belonging to tissue $t$. The z-based TS score measures the distance from the tissue median to the population mean standardized by the population standard deviation. It standardizes gene expression across genes and tissues, making expressions comparable.

**Protein TS score filtering**

If a tissue only has one replicate, NA is assigned and its score will be marked as “NA_one_rep_in_raw”. If a gene is only observed in one experimental run, we marked its tissue scores as “NA_only_form_one_run”. The protein TS scores and z-scores based on sample median and MAD are summarized in Table S3 with filter information.

**Protein enrichment category**

We defined four enrichment categories based on protein TS scores: tissue-specific, tissue-enriched-but-not-specific, house-keeping and others. We required the genes non-NAs in at least three tissues in their TS scores in the categories of tissue-specific and tissue-enriched-but-not-specific. If there is at least one TS score $\geq$ 4 and there are no TS scores lying in the interval (2.5, 4), we called this protein tissue specific (in total 1586). If there is at least one TS score $\geq$ 2.5 but the protein is not tissue specific, we called this protein tissue enriched but not specific (in total 3921). If in the raw abundance expressions, each tissue is observed in at least one replicate and all its TS scores are less than 2 and no missing values in its scores, we called this protein house-keeping (in total 1578). The rest proteins belong to the “others” category, in total 5542 genes. The enrichment category comparison with RNA is summarized in Table S6.

**3.2 Tissue specificity score in RNA expression**

**RNA population**

We transformed the TPM in log scale for our RNA expression analysis. After we adjusted the near-zero-TPM by a small value, the distribution of TPM in log scale is smoothened. Similar to our protein expression analysis, when comparing samples from multiple tissues, the main effect in the population in RNA expression is the tissue effect, which is confirmed from the hierarchical cluster plot as shown in Fig.S4. As an example, Fig.S11 is the data from 16 genes, each gene has one big density bump and the outliers are located in the tails. Several previous studies assumed the Gaussian noise when analyzing log transformed RNA expression (*14, 15*). In this study, we applied our population fitting algorithm (AdaTiSS) on each gene. Fig.S12 shows the histogram of the fitted population in RNA using our algorithm. All the population fitting parameters in RNA can be found in Table S2.

In addition, we compared our results to the standard sample median and sample MAD. Fig.S13 shows the gene-wise comparisons for the population mean and population standard deviation (SD) on 12,238 protein-and-RNA commonly quantified genes. We found there are 48 genes that have fitted SD/MAD ratios that are greater than 1.2 and 1,392 genes have a ratio that is less than 0.8. Examples of these fittings can be found in Fig.S14-S16. We provide all the population information and population fitting for RNA on our website.

**RNA TS scores**

The same definition as protein TS scores.

**RNA TS score filtering**

For genes with low expressions, we assigned the TS scores as NAs and marked them as “NA_all_tissues_tpm_less_1”. We filtered the TS score as NAs in RNA if such tissue has score greater than 2.5 but its raw sample median TPM less than 1 and marked it as “NA_raw_tpm_less_1”. The RNA TS scores and the z-scores based on sample median and MAD are summarized in Table S2 with filtering information.

**RNA enrichment category**

The same definition as the protein enrichment category. Among 12,238 protein and RNA commonly quantified genes, there are 1,097 tissue-specific genes, 5336 tissue-enriched-but-not-specific genes, 2595 house-keeping genes.

**3.3 Tissue specificity score comparison in protein and RNA**

We compared protein and RNA TS scores in two levels: (i) across tissues and (ii) in single tissue level.

Across tissues, we did the Spearman correlation between the protein TS scores and RNA TS scores for each gene. For the analysis, at least five non-NAs tissues in both protein and RNA were required. The p-values are calculated from permutation test based on200 permutations and are adjusted from the Benjamini-Hochberg procedure (*16*). The results are summarized in Table S3. 10,317 genes were compared and the false discovery rate (FDR) was controlled for at ≤ 0.05. We found there were 6,604 genes, protein-and-RNA significantly positively correlated and 92 protein-and-RNA significantly negatively correlated, and the remaining 3,621 protein and RNA were non-significantly correlated.

We jointly defined outliers in a (RNA, protein) 2-dimensional analysis and gave configuration of outlying tissues. We considered the genes that commonly were quantified in both RNA and protein from the mapped samples. We extended the work in AdaRTeg and AdaTiSS to two-dimensional fitting and robustly fit the population information for the genes observed in at least 50 samples that are from at least 10 tissues in both RNA and protein. For the rest of the genes, we forced the protein and RNA sample population correlation to be zero. Based on the marginal robust z-scores, we defined the outlier region under Mahalanobis distance for gene $i$ by

$$R_{i}=\left\{ z_{i}\in\mathbb{R}^{2}:\sqrt{z_{i}^{\top}\hat{\Sigma}_{i}z_{i}}\geq2.5 \right\}\text{where}\text{ }\hat{\Sigma}_{i}=\left( \begin{matrix} 1 & \hat{\rho} \\ \hat{\rho} & 1 \end{matrix} \right),$$

and called the tissue an outlier if its sample joint z-scores

$$Z_{ij}^{(joint)}=(Z_{ij}^{(RNA)}, Z_{ij}^{(prt)})$$

satisfying

$$\#\{Z_{ij}^{\left( joint \right)}\in R_{j}\}>1 and sample proportion of \{Z_{ij}^{(joint)}\in R_{j}\}\geq50\%.$$

The tissue joint specificity score was defined by $S_{it}^{(joint)}=(S_{it}^{(RNA)}, S_{it}^{(prt)})$ where S is defined in a similar way as in one dimension. Based on $S_{i.}^{(joint)}$’s, we categorized RNA-protein outlying configurations and assigned each outlier tissue a category based on the maximum of the cosine similarities between the joint TS score and 8 directions: (1) $(0,1)$ direction — only protein are highly expressed, (2) (1,1) direction — both protein and RNA are highly expressed, (3) (1, 0) direction — only RNA are highly expressed, (4) (1,-1) direction — protein low expressed but RNA highly expressed, (5) (0,-1) only protein are low expressed, (6) (-1,-1) both protein and RNA are lowly expressed, (7) (-1,0) only RNA lowly expressed, (8) (-1, 1) protein highly expressed but RNA lowly expressed.

In a single tissue level, we defined two concepts of concordance and discordance. If a gene has TS scores in both RNA and protein greater than 2.5 in a certain tissue, it is concordant in that tissue. If a gene has a protein (RNA) TS score greater than 2.5 and its protein (RNA) score is at least 1.5 units higher than its RNA (protein) score, the gene is discordant in that tissue. We summarized the tissue concordance and discordance results in Table S3.

**3.4 Tissue specificity score in isoforms**

We applied the same procedures of obtaining protein and RNA TS scores in the gene level for the isoform level. The protein and RNA isoform TS scores and their enrichment comparison results are summarized in Table S5.

**4. Methods in the analysis**

**Gene enrichment analysis**

We did enrichment analysis of Kyoto Encyclopedia of Genes and Genomes (KEGG) and Gene Ontology (GO) (*17*) terms based on R package “STRINGdb” (*18*). The GO term enrichment results for tissue-enriched proteins in individual tissues and the results for house-keeping proteins are all summarized in Table S3. The pathway enrichment results for tissue-enriched proteins in individual tissues are summarized in Table S3. For the enrichment results in the metabolism pathway, we obtained its sub-pathway IDs from KEGG under “Metabolism” and summarized the results in Table S3.

**Enrichment analysis of unidentified proteins**

The Human Protein Atlas project (HPA) annotated protein classes for 19,628 genes (*19*). There are 12,909 genes in our data mapped to the list. We applied Fisher’s exact test to see whether the unidentified proteins are enriched in a certain predicted protein class. The results (summarized in Table S7) showed that the unidentified proteins are significantly enriched in the membrane protein class, but not significantly enriched in the intercellular nor the secreted protein class.

**Association analysis of protein identification and RNA expression level**

We investigated whether a gene that is identified in protein is associated with its RNA expression abundance. In our analysis, we only considered the genes that have RNA expressions at least in one tissue with a raw TPM > 1. In total, there are 17,690 such genes. The mean RNA expression level (in log scale) across 32 tissues was used as the gene’s RNA level expression. The RNA expression abundances are grouped into 10 bins: $(-\infty,1],(1,2], \ldots, (8,9], (9,+\infty)$. We applied a chi-square test for independence of gene expression in RNA and protein identification. We found that when we tested on all the bins, the p-value is less than $2.2\times{10}^{-16}$, but if we only tested on the bins starting from $(5,6]$, the p-value is 0.14, which indicates when RNA average expression is higher than 32 in raw TPM, the identification of the corresponding protein is not significantly associated with its RNA expression. The contingency table is summarized in Table S8.

**Association analysis of protein isoform identification and RNA isoform expression level**

Similarly as in the gene level, we investigated whether the identification of protein isoform is associated with RNA abundances. Genes having at least two isoforms in RNA are included in the study. The mean of RNA expressions (in log scale) from 32 tissues is used as RNA isoform level expression. The RNA isoform expressions are grouped into six bins: $(-\infty,1],(1,2], (2,3], (3,4], (4,5], (5,+\infty)$. We counted the number of rank 1 RNA isoforms with protein evidence in each expression bin and summarized the information in Table S9(a). In addition, the same analysis was performed for the rank 2 RNA isoforms and the information is summarized in Table S9(b). We applied chi-square test for testing independence of RNA isoform expression and protein identification. For the rank one isoforms, protein identification is not significantly associated with the RNA expression under significance level of 0.05, while for the rank two isoforms, RNA isoform expression may affect the identification of the protein isoform.

**Tissue enriched proteins and genetic diseases**

In this analysis, we investigated the link between the tissue enriched proteins and genetic diseases. The enriched proteins in each tissue were compared to the OMIM disease gene list (*20*) with disease-relevant tissue abnormality. Fisher’s exact test was applied to assess the significance of the overlap. The results are summarized in Table S11.

**Single nucleotide polymorphisms variant (SNV) in protein**

All the spectra were searched against the GENCODE V19 database populated with SNV peptides at peptide FDR of 1%. The SNV peptides were generated based on the genomic information available from 12 individuals in our study. We also did a de no vo search of SNV peptides using PEAKS Studio. Only the overlapped SNP peptides were kept for further analysis. To improve the confidence level, we added another layer of test to verify if the SNV peptides were identified in the expected sample (individual) based on the known genetic information. Since SNV candidate peptides have high missing values, we took sample median + 2.5 MAD as the threshold based on the observed samples and the outliers were the ones whose abundances exceed the threshold. Then we applied Fisher’s exact test for testing the enrichment of SNV peptides in the correct samples. With controlled FDR under level 0.1, there are 127 significantly enriched SNV peptides. The results are summarized in Table S13.

**Supplementary Figures**

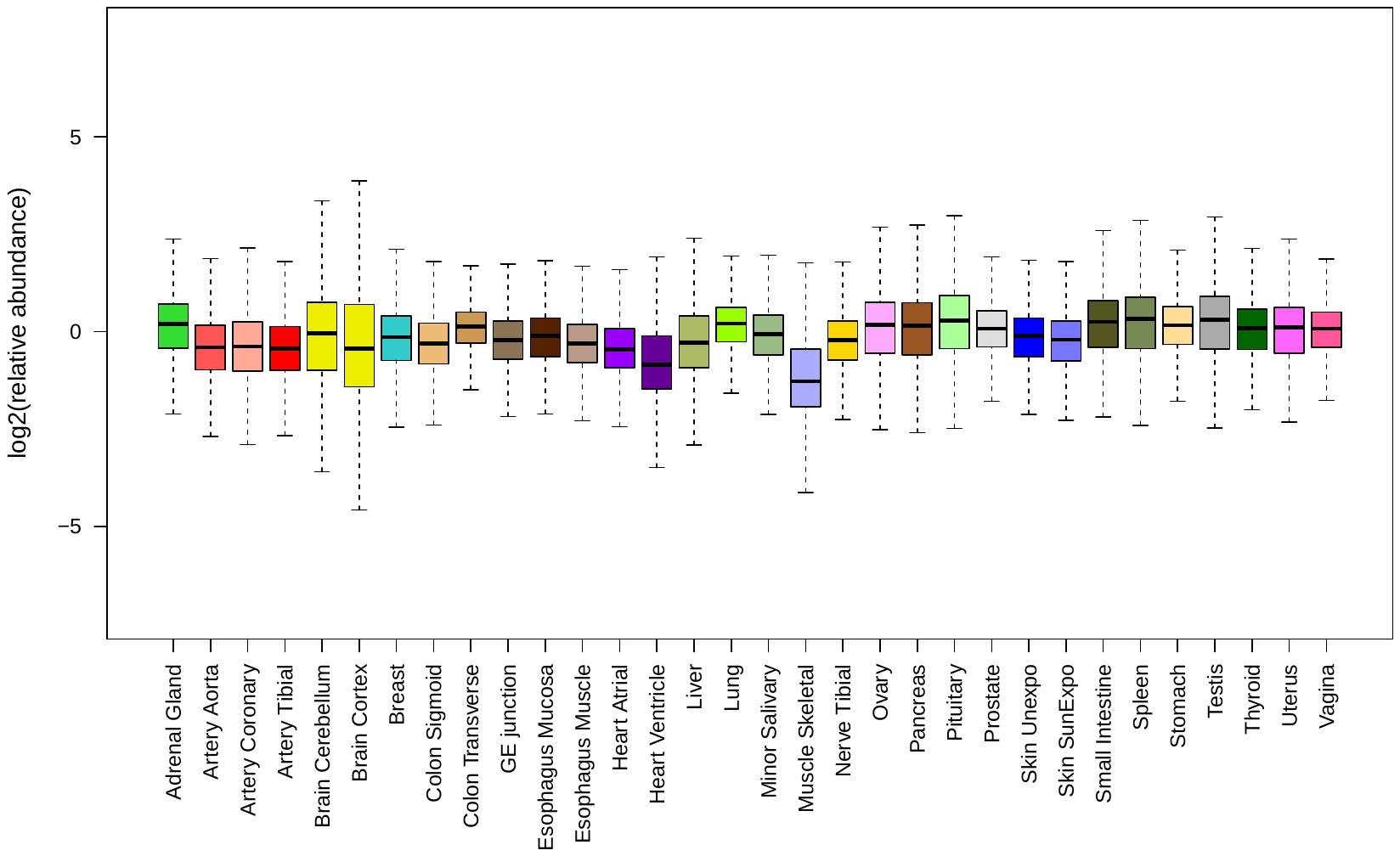

**a**

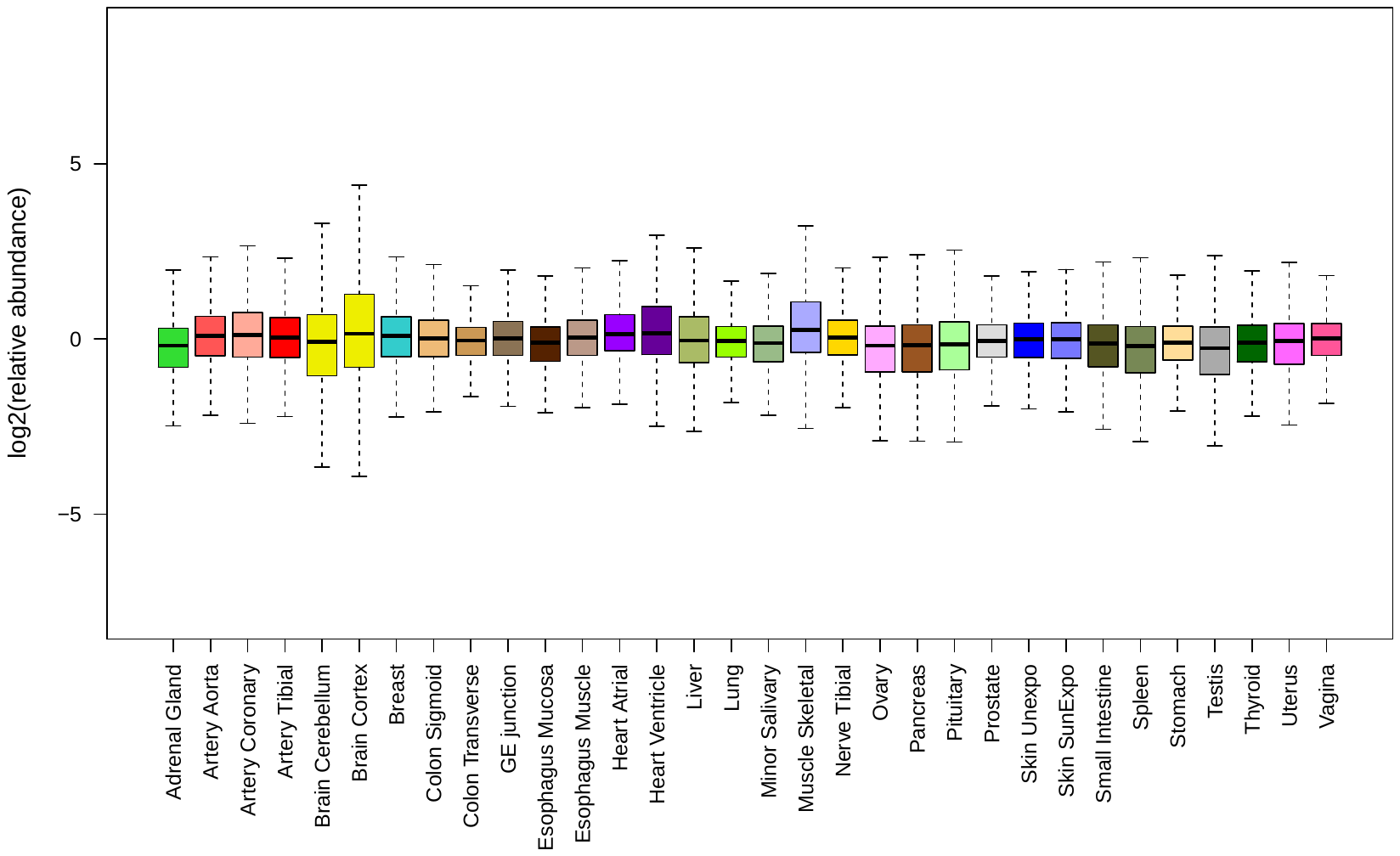

**b**

**Fig.S1: Boxplots of tissue sample medians in protein profile before robust normalization (a) and after robust normalization (b).**

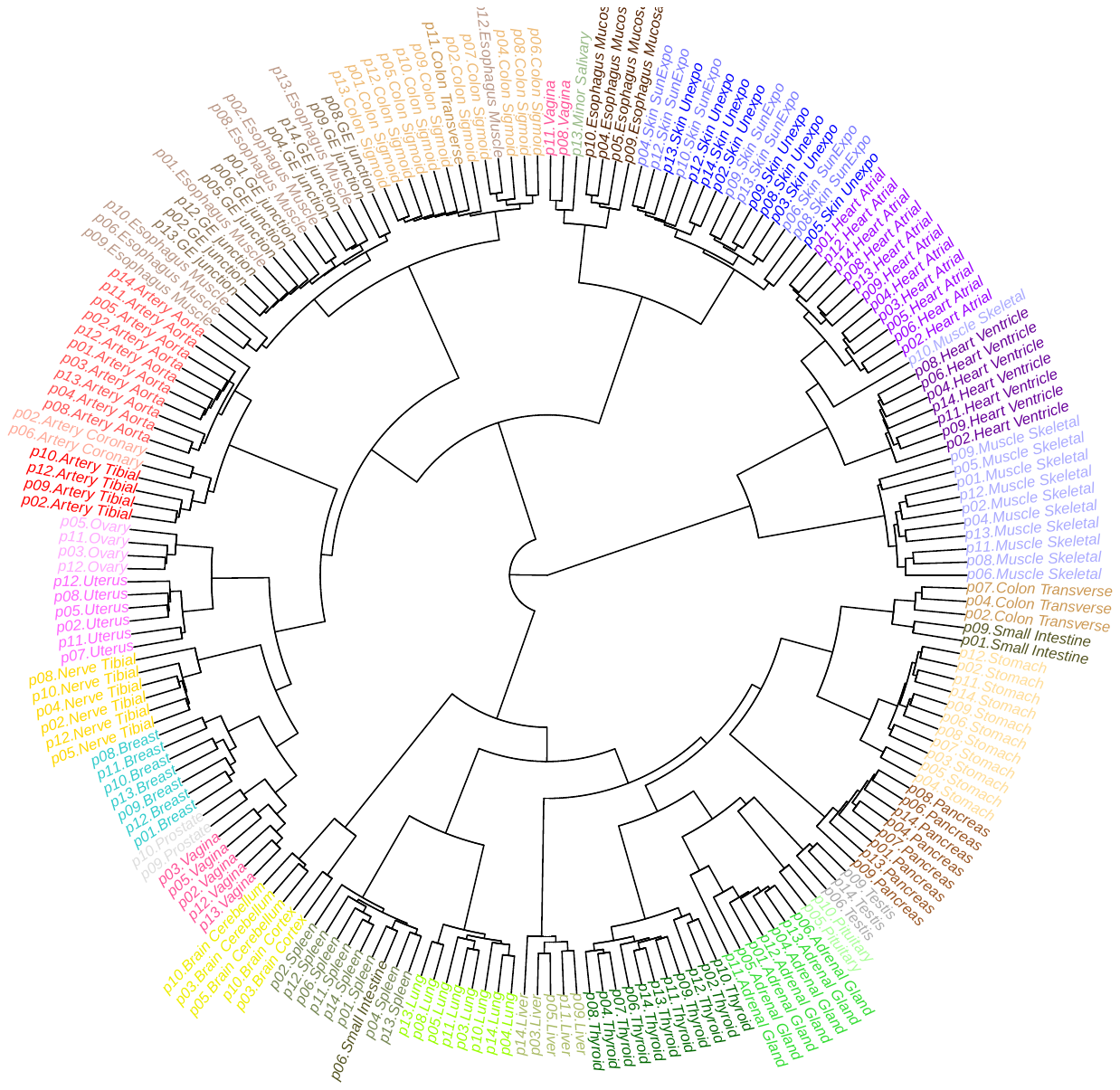

**Fig.S2: Dendrogram of tissue samples in protein expression based on pairwise Euclidean distance from Ward’s method.** The sample names are labeled by the people index p01 – p14 followed by the tissue name.

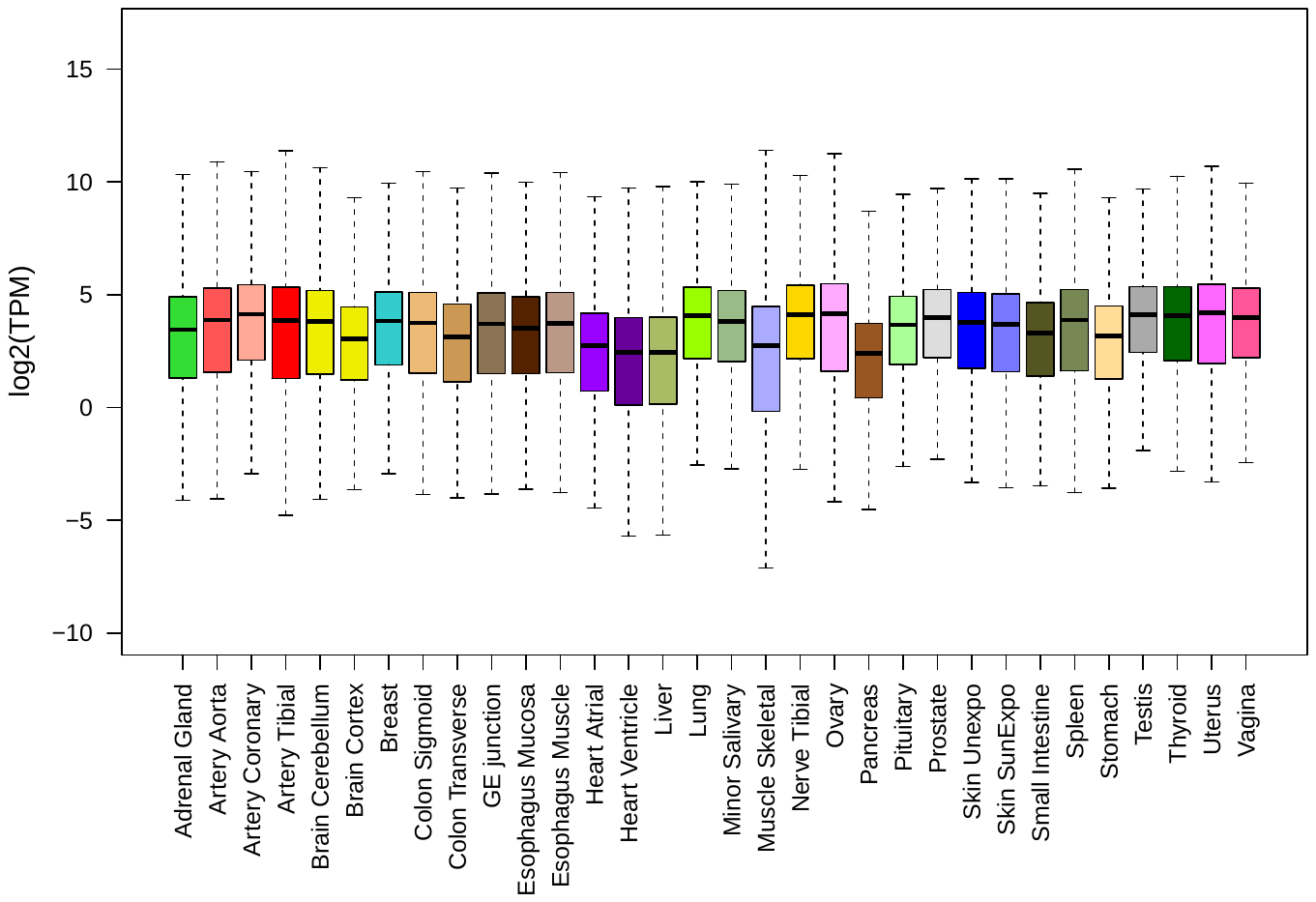

**a**

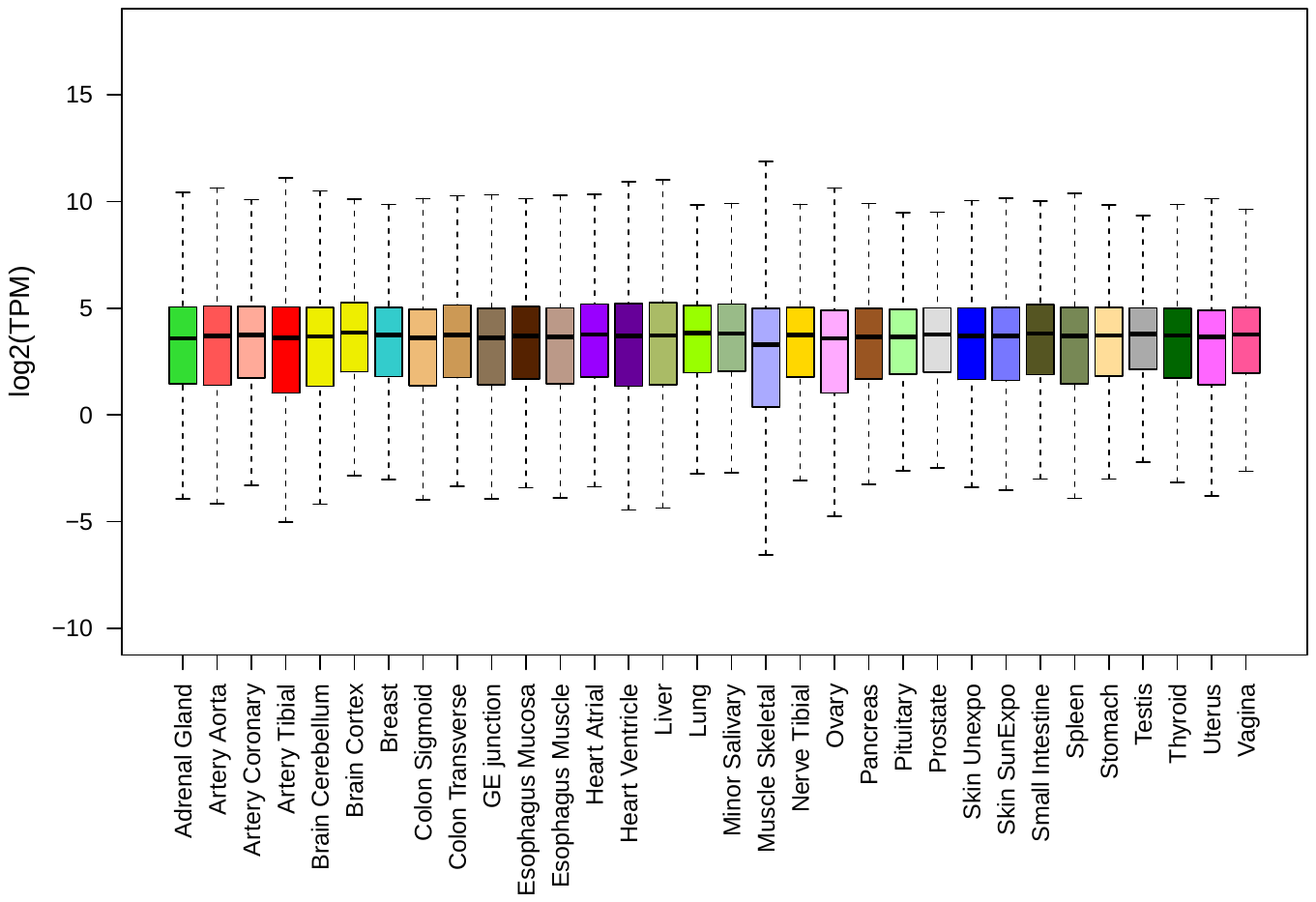

**b**

**Fig.S3: Boxplots of tissue sample medians in RNA profile from protein corresponding samples and commonly quantified genes before robust normalization (a) and after robust normalization (b).**

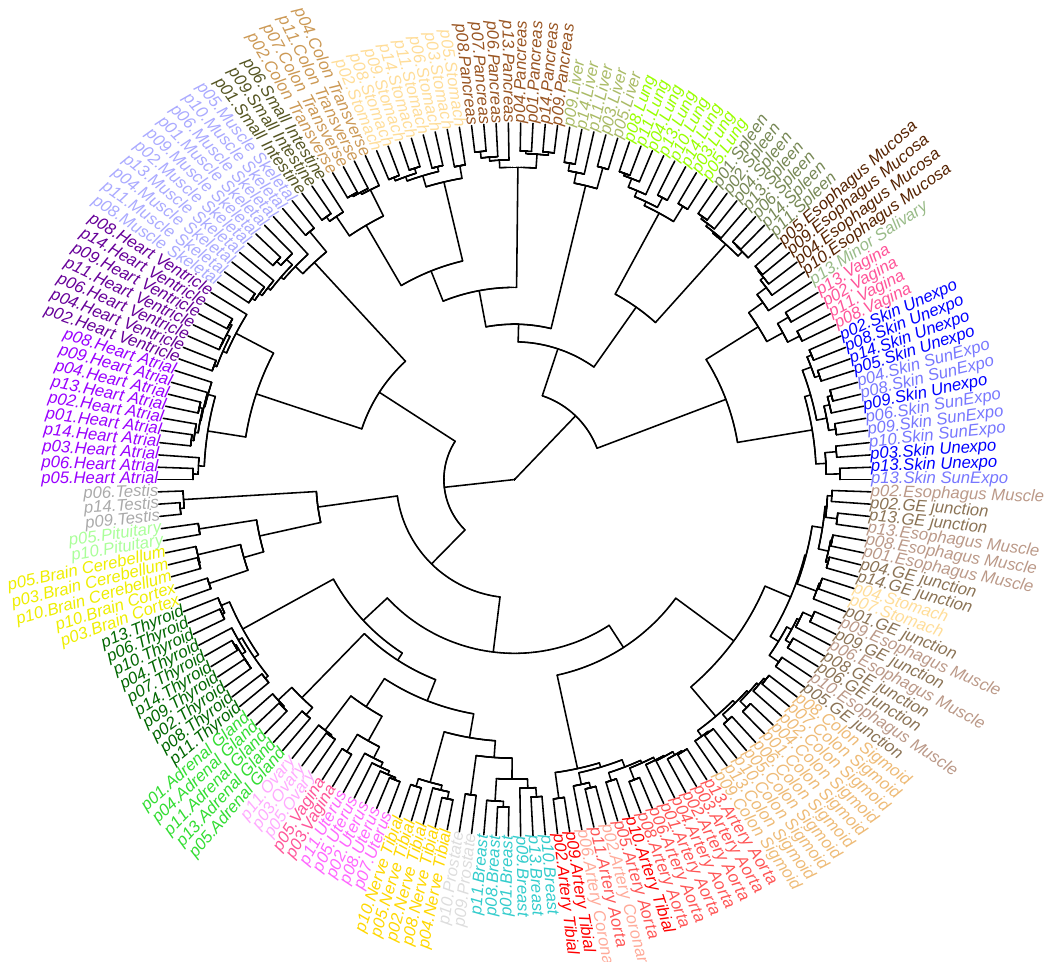

**Fig.S4: Dendrogram of tissue samples in RNA expression of RNA and protein commonly quantified genes based on pairwise Euclidean distance from Ward’s method.** The sample names are labeled by the people index p01 – p14 followed by the tissue name.

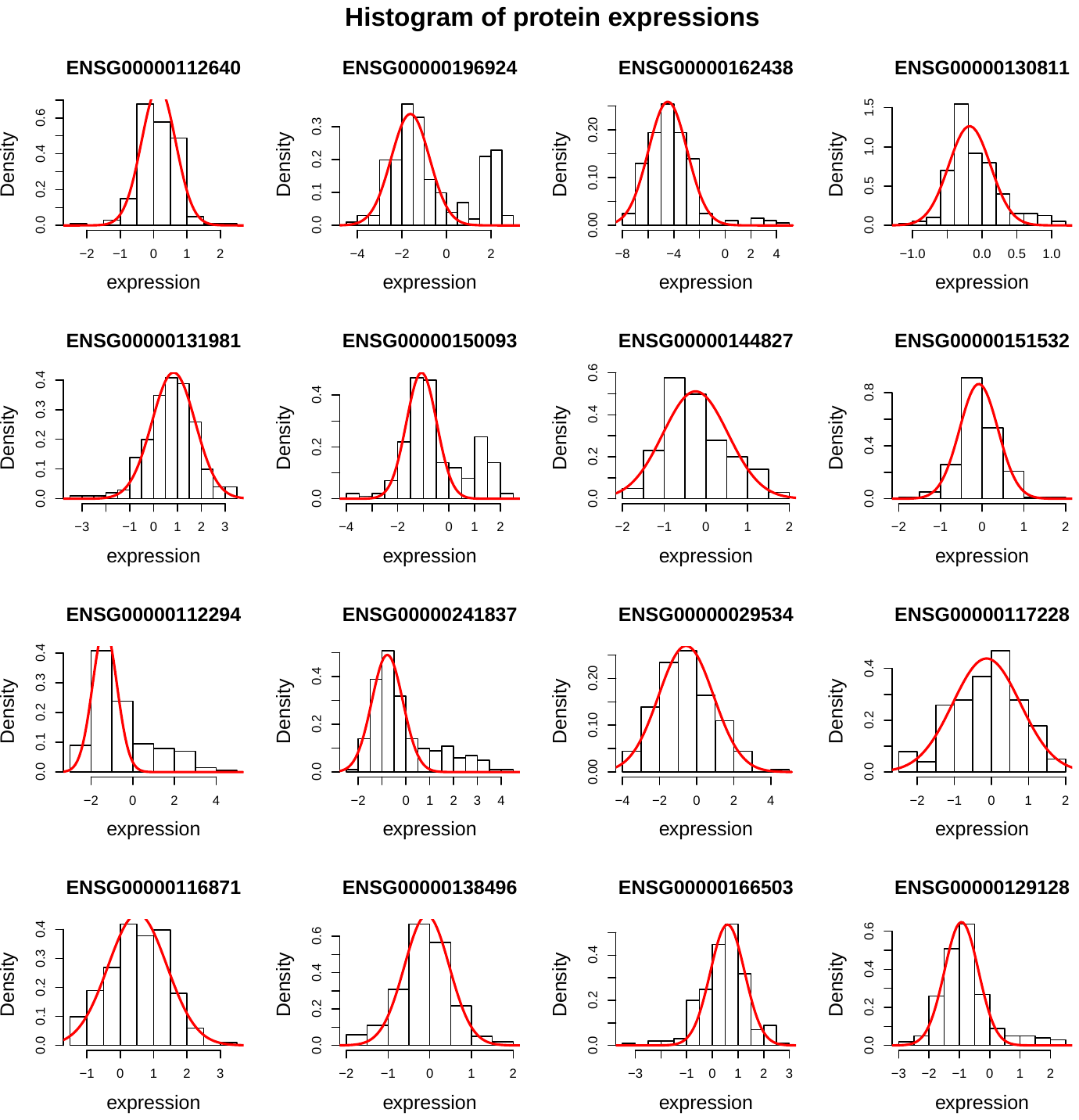

**Fig.S5. Histograms of protein expressions from randomly selected 16 genes without missing values.** Each histogram is for one gene. The red curve is from our robustly fitted population density. There is a big density bump for each gene shown here.

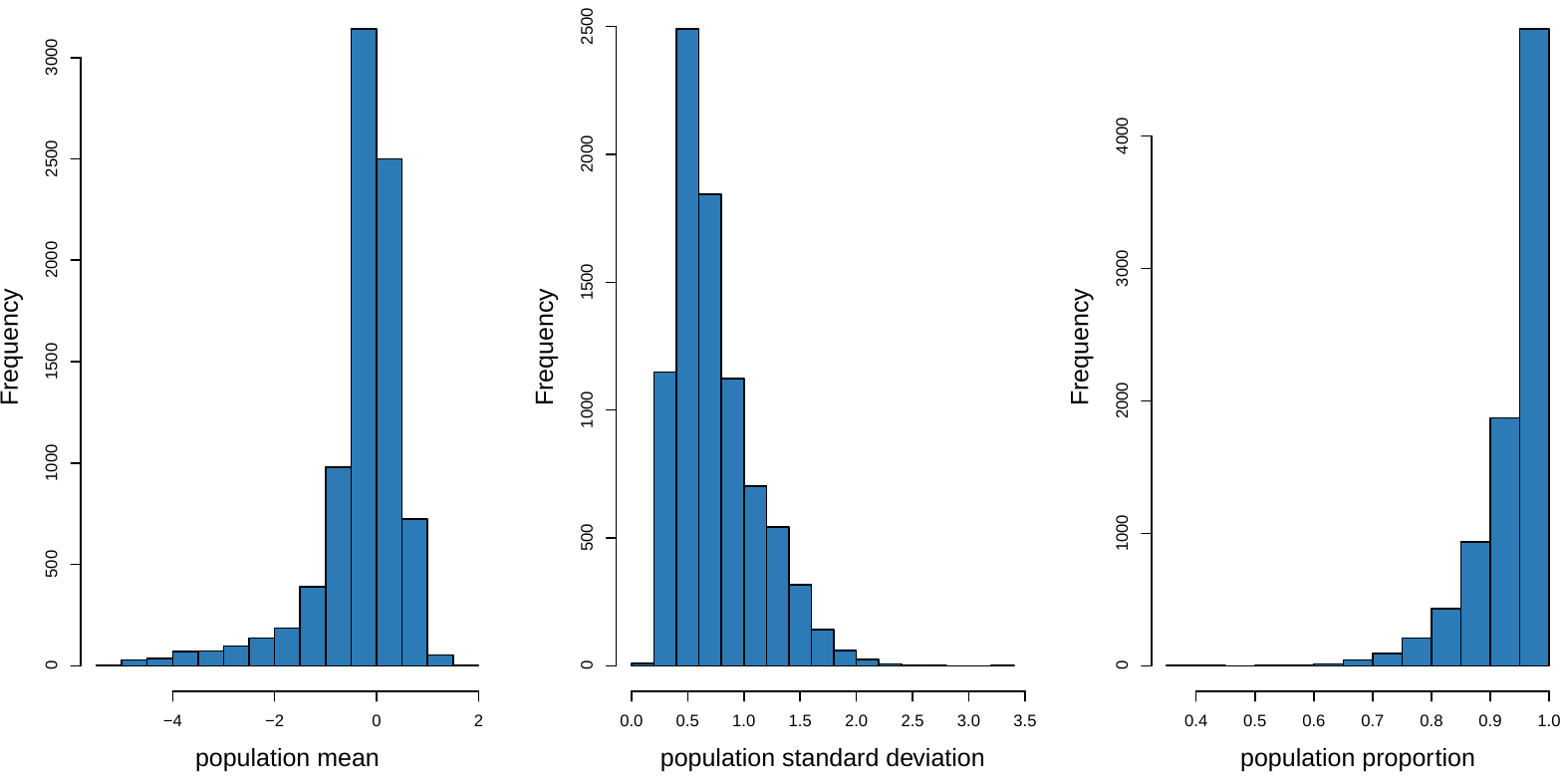

c

b

a

**Fig.S6. Histograms of protein population parameters fitted from our algorithm AdaTiSS.** Panel a shows this histogram for population mean; Panel b for population standard deviation and Panel c for population proportion (due to fitting approximation, truncated by 1).

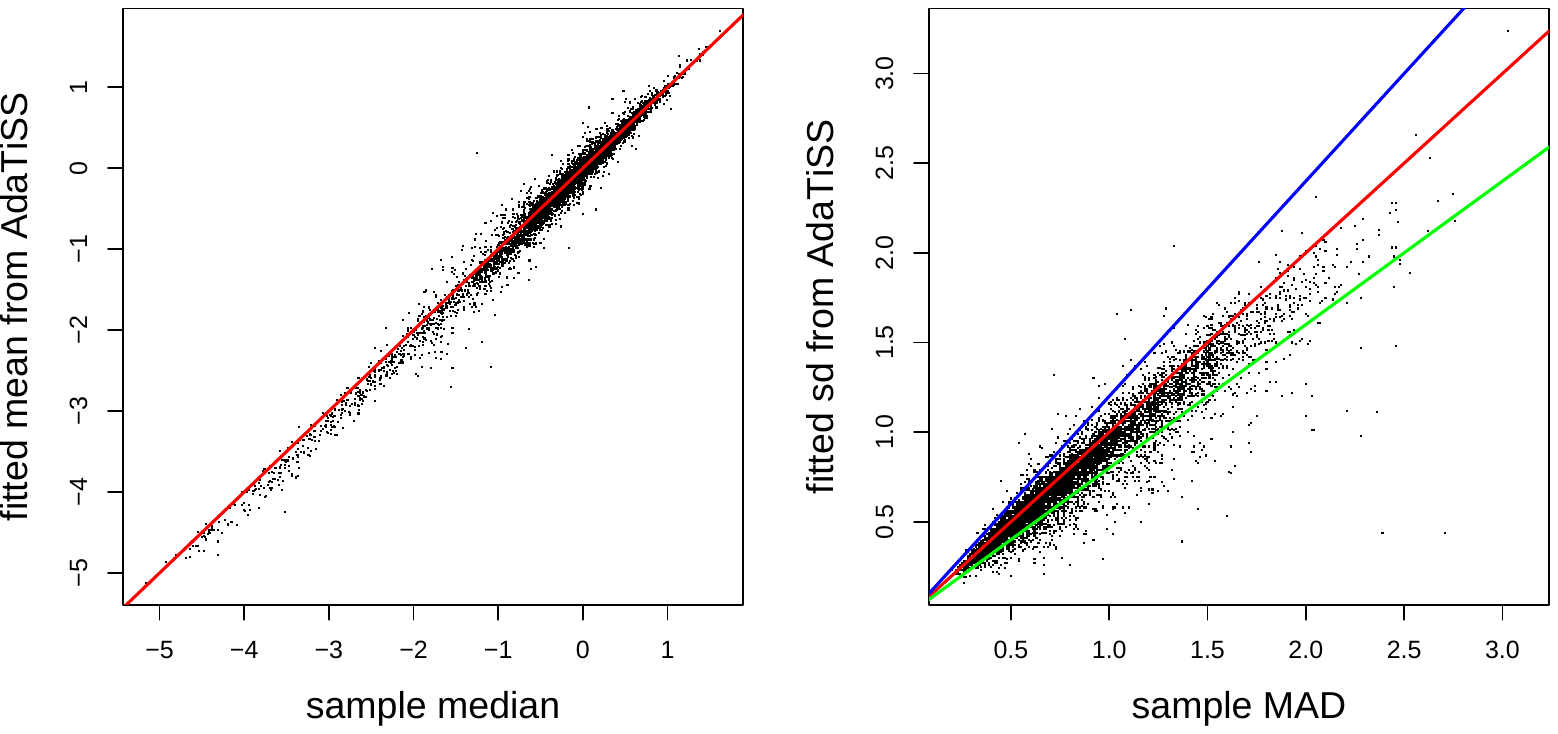

b

a

**Fig.S7. In protein, gene-wise comparison between sample median vs. fitted mean from AdaTiSS for estimating population mean in Panel (a) and between sample MAD vs. fitted mean from AdaTiSS for estimating population standard deviation from AdaTiSS in Panel (b)**. One dot is for one gene. The red line is the identity line y=x. The blue and green lines in Panel (b) are for y=1.2x and y=0.8x respectively.

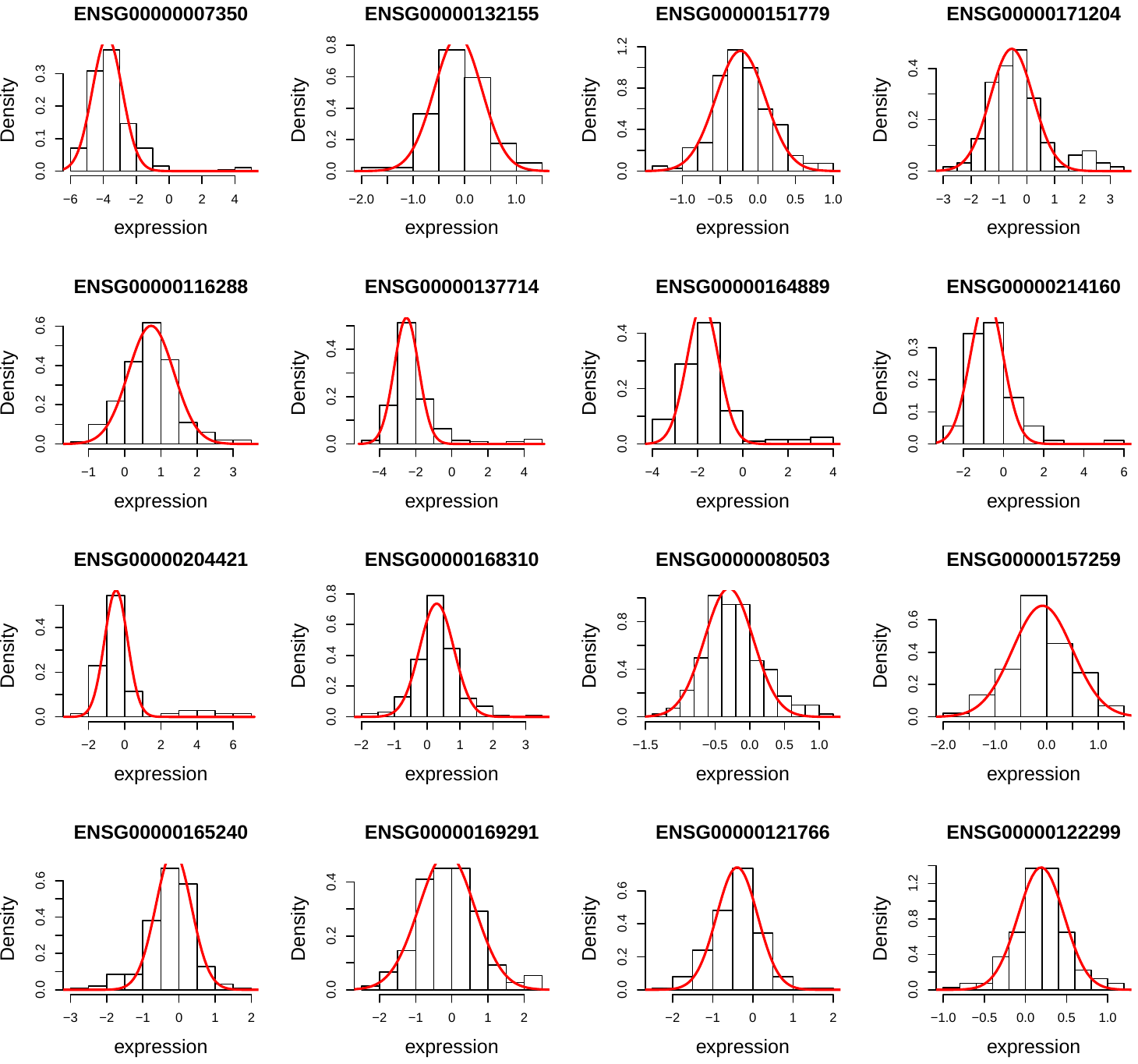

**Fig.S8. Histograms of protein expressions in randomly selected genes having similar MAD and fitted standard deviation from AdaTiSS.** The red curve is the density fitting from AdaTiSS. For these genes, the ratio of fitted standard deviation to MAD is between 0.8 and 1.2.

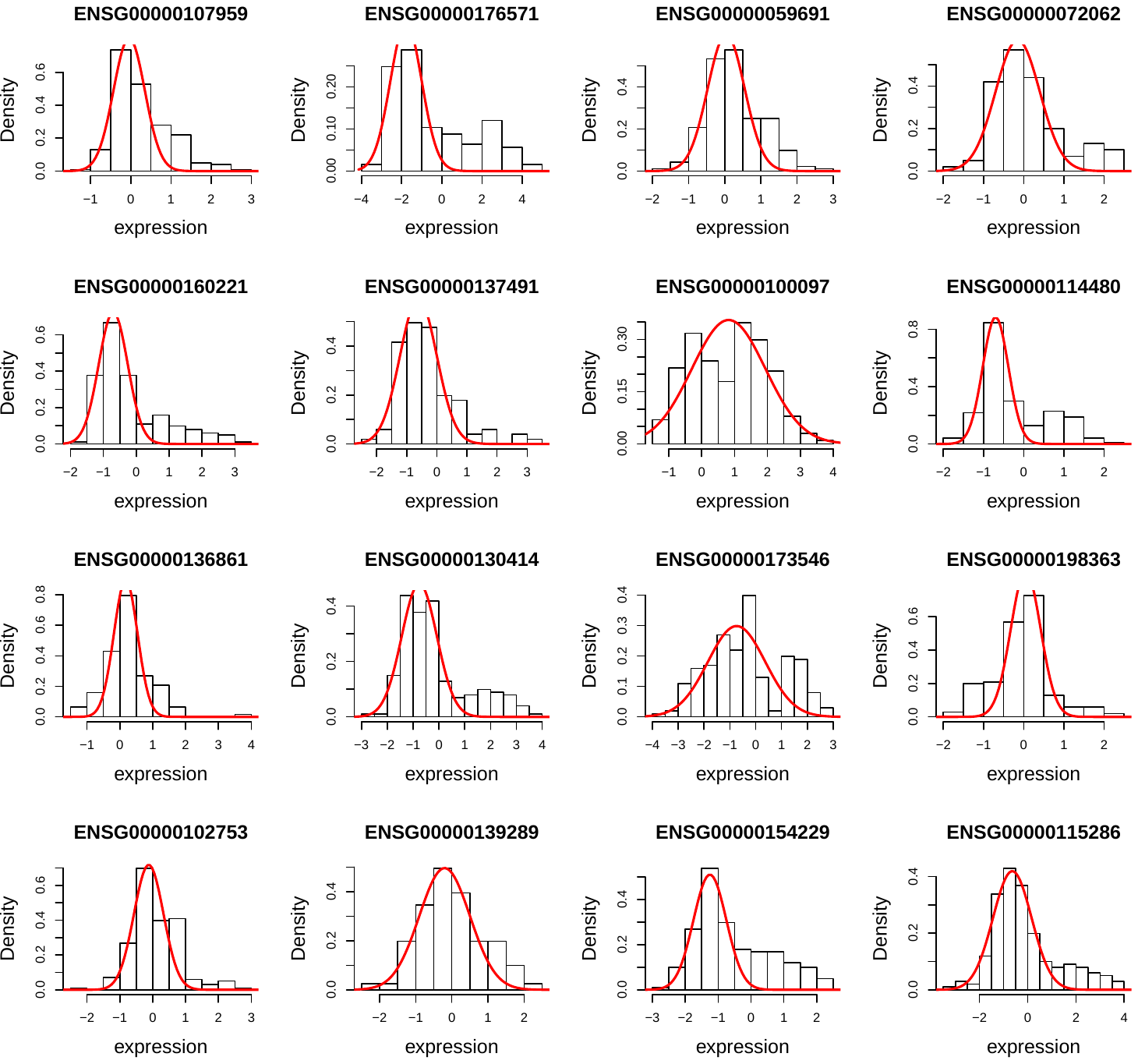

**Fig.S9. Histograms of protein expressions in randomly selected genes whose ratio of and fitted standard deviation from AdaTiSS to MAD < 0.8.** The red curve is the density fitting from AdaTiSS.

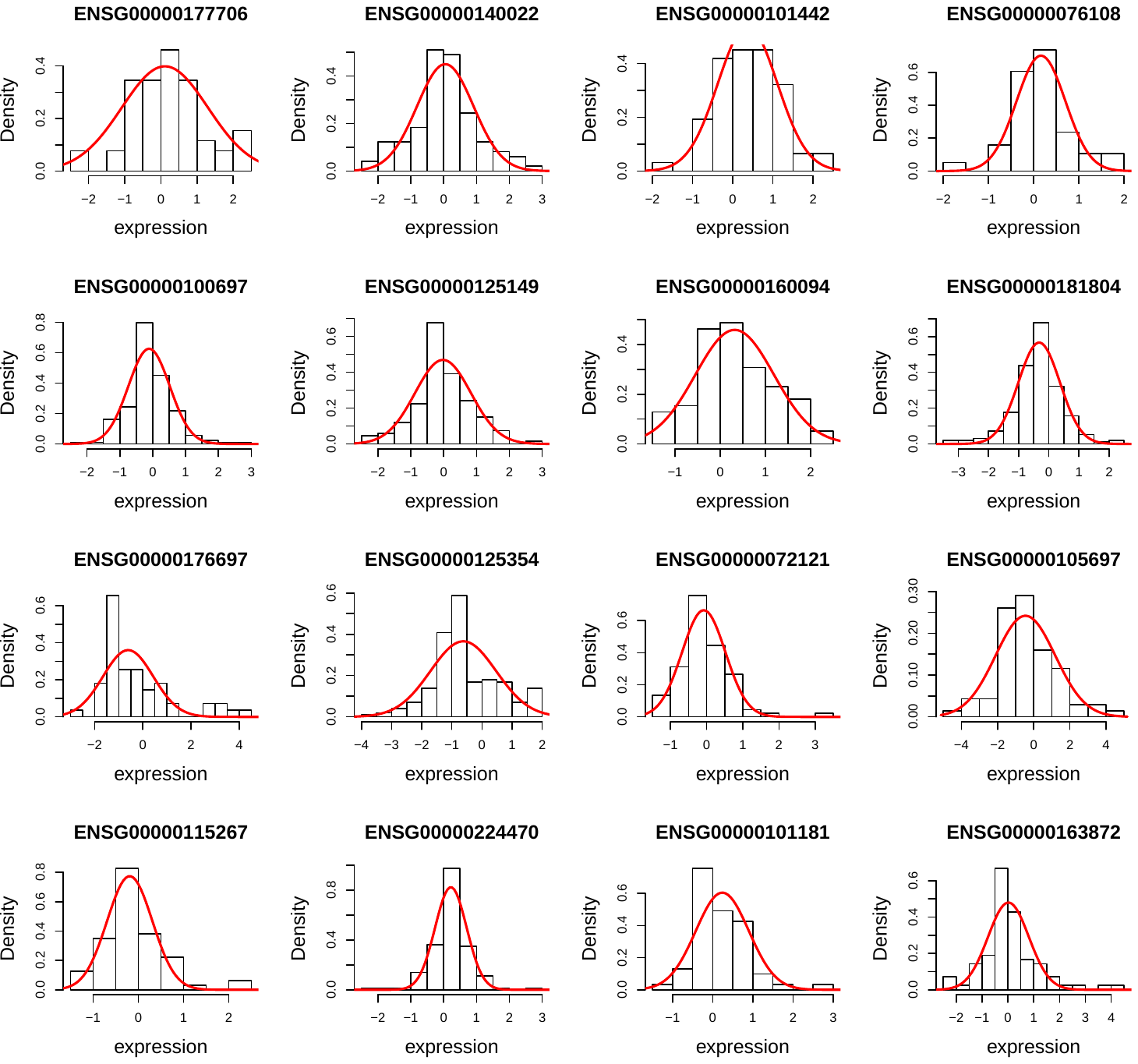

**Fig.S10. Histograms of protein expressions in randomly selected genes whose ratio of and fitted standard deviation from AdaTiSS` to MAD > 1.2.** The red curve is the density fitting from AdaTiSS.

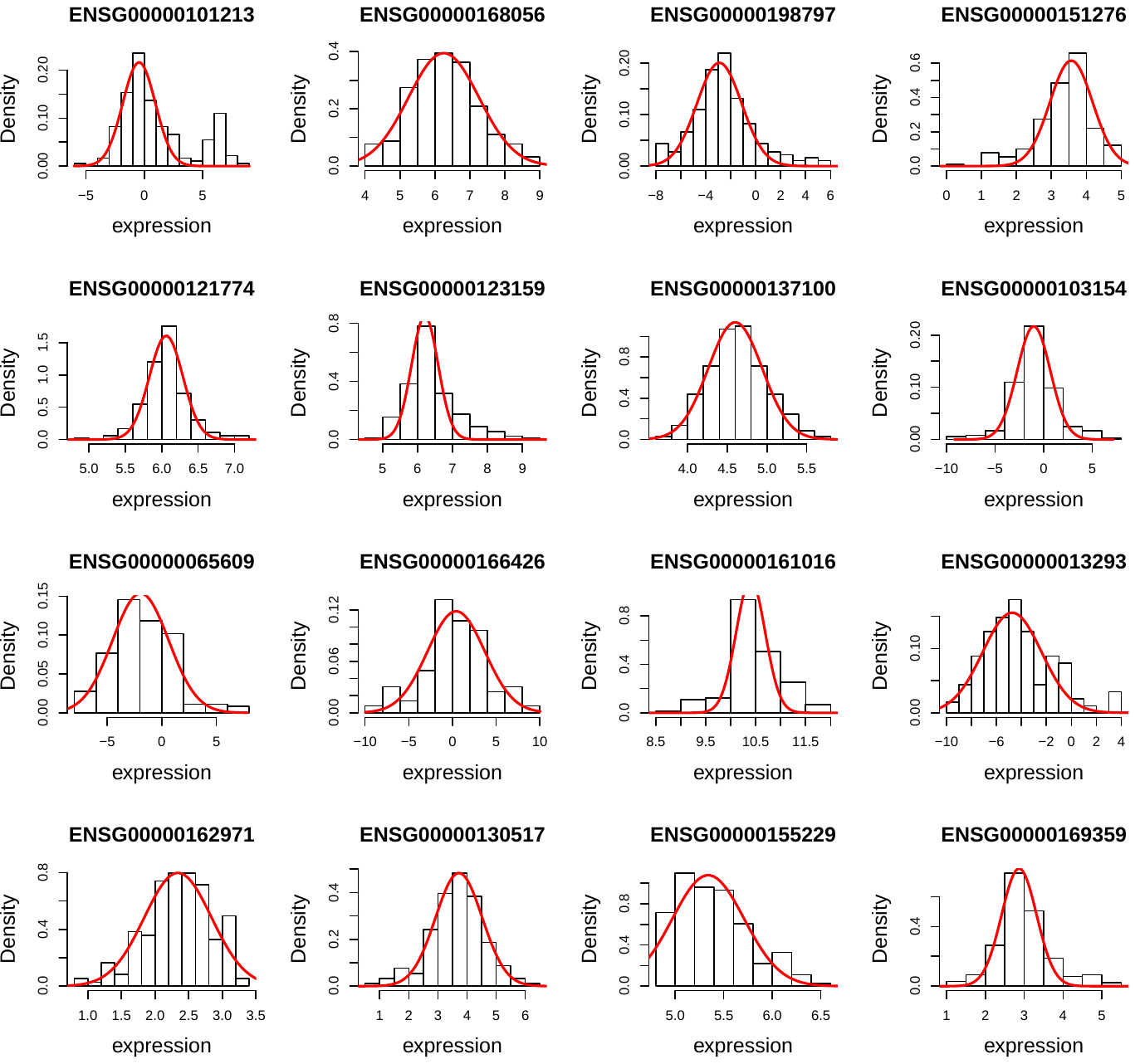

**Fig.S11. Histograms of RNA expressions from randomly selected 16 gene.** Each histogram is for one gene. The red curve is from our robustly fitted population density. There is a big density bump for each gene shown here.

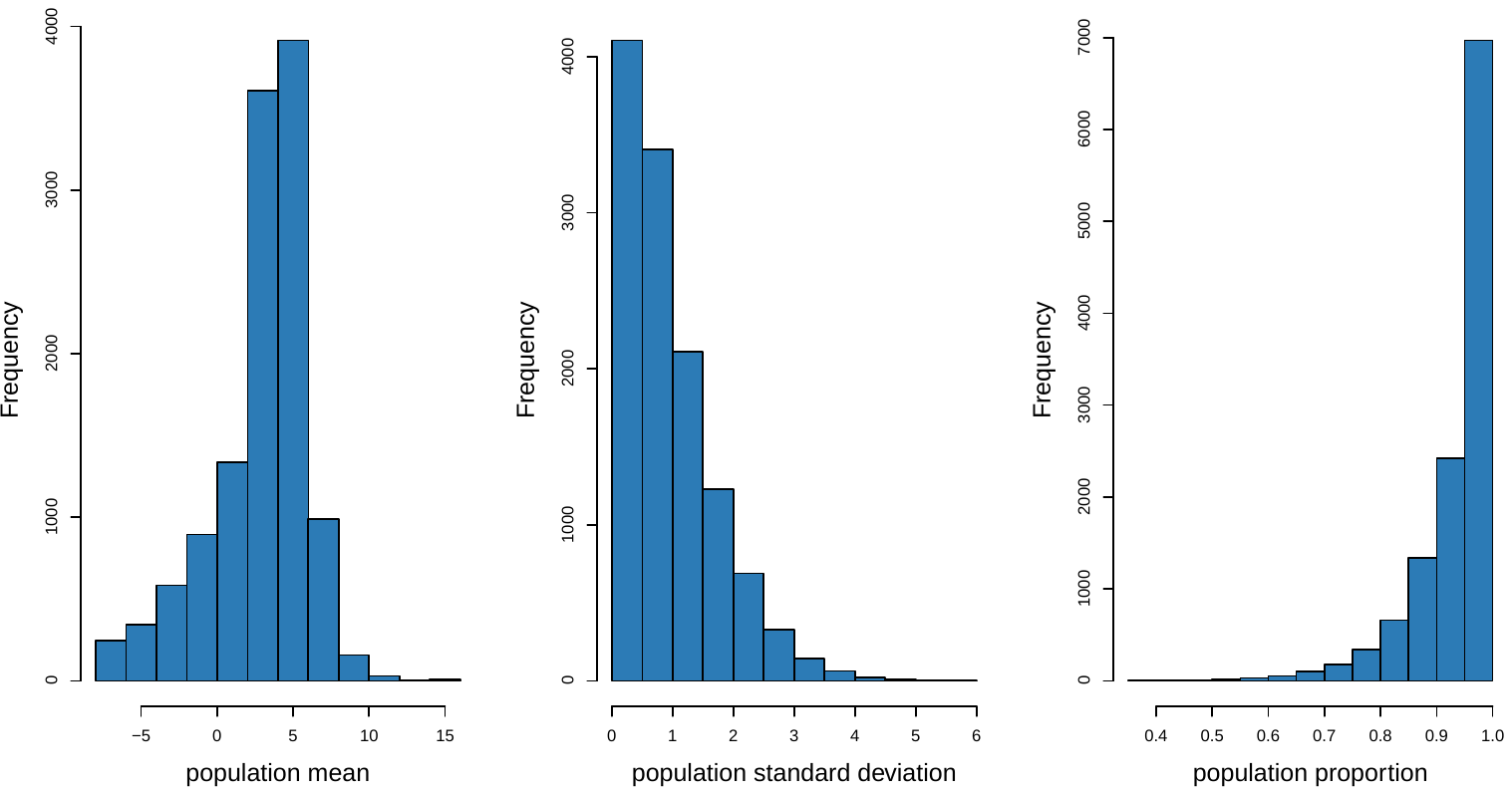

c

b

a **Figure S7. Gene-wise comparison between sample median vs. fitted mean from AdaTiSS for estimating population mean in Panel (a) and between sample MAD vs. fitted mean from AdaTiSS for estimating population standard deviation from AdaTiSS in Panel (b)**. One dot is for one gene. The red line is the identity line y=x. The blue and green lines in Panel (b) are for y=1.2x and y=0.8x respectively.

**Fig.S12. Histograms of RNA population parameters fitted from our algorithm AdaTiSS.** Panel (a) shows this histogram for population mean; Panel (b) for population standard deviation and Panel (c) for population proportion (due to fitting approximation, truncated by 1).

b

a

**Fig.S13. In RNA, gene-wise comparison between sample median vs. fitted mean from AdaTiSS for estimating population mean in Panel (a) and between sample MAD vs. fitted mean from AdaTiSS for estimating population standard deviation from AdaTiSS in Panel (b)**. One dot is for one gene. The red line is the identity line y=x. The blue and green lines in Panel (b) are for y=1.2x and y=0.8x respectively.
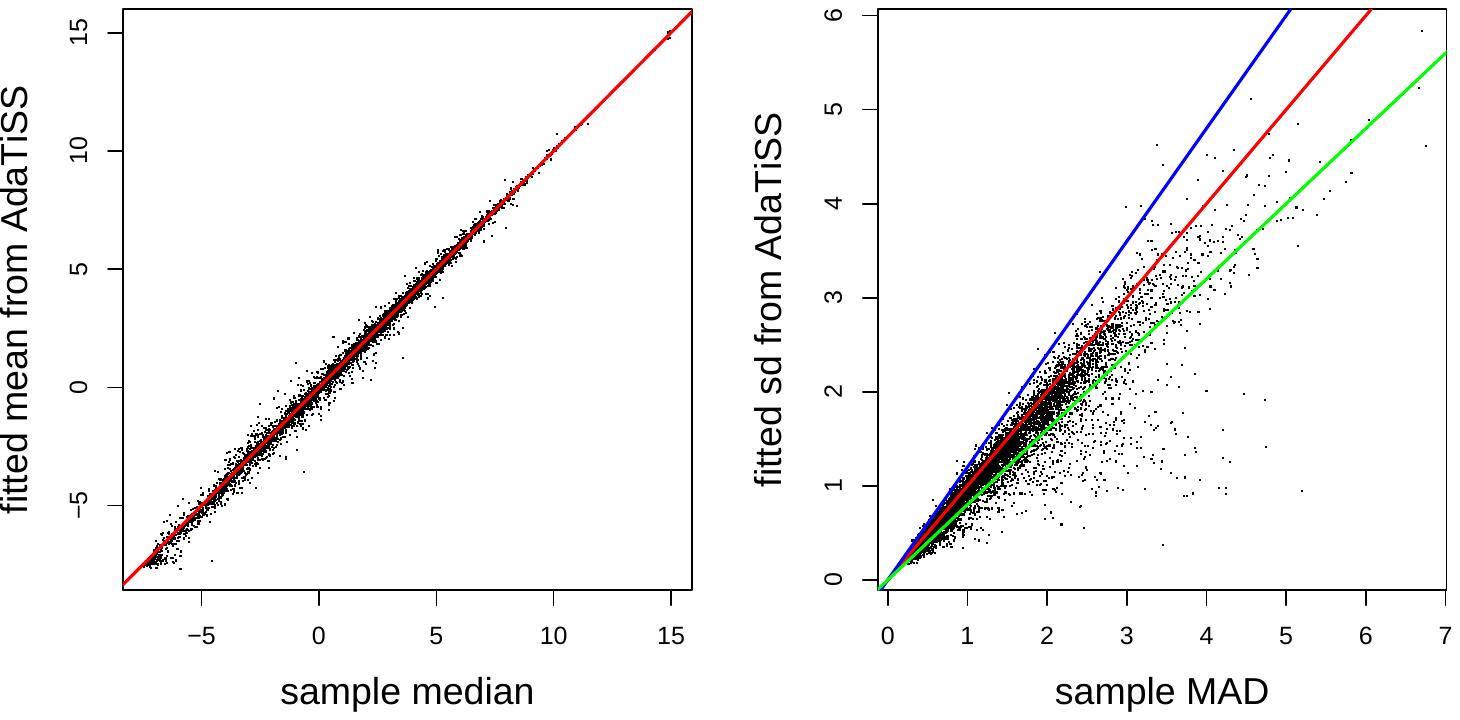

**Fig.S14. Histograms of RNA expressions in randomly selected genes having similar MAD and fitted standard deviation from AdaTiSS.** The red curve is the density fitting from AdaTiSS. For these genes, the ratio of fitted standard deviation to MAD is between 0.8 and 1.2.
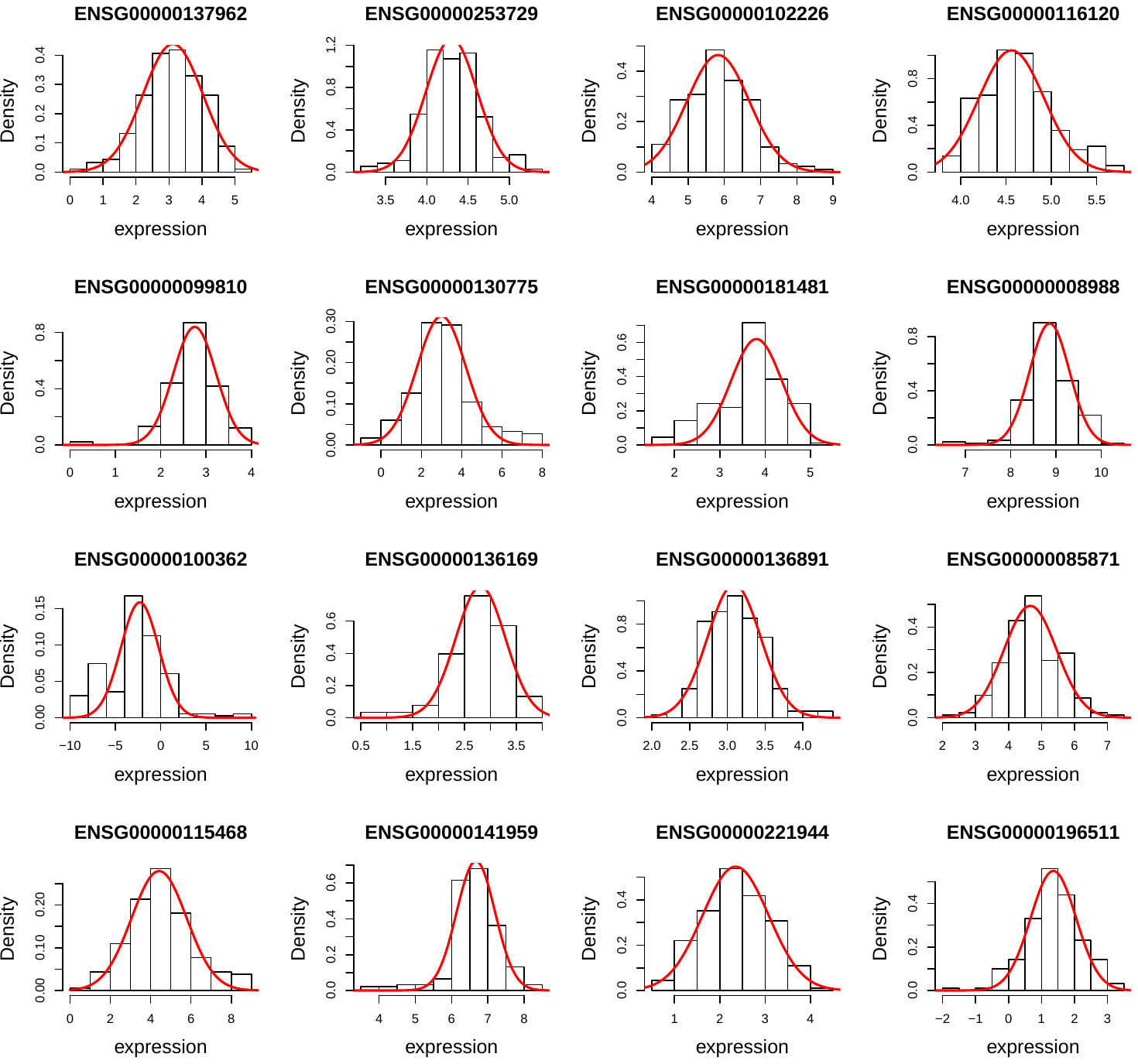

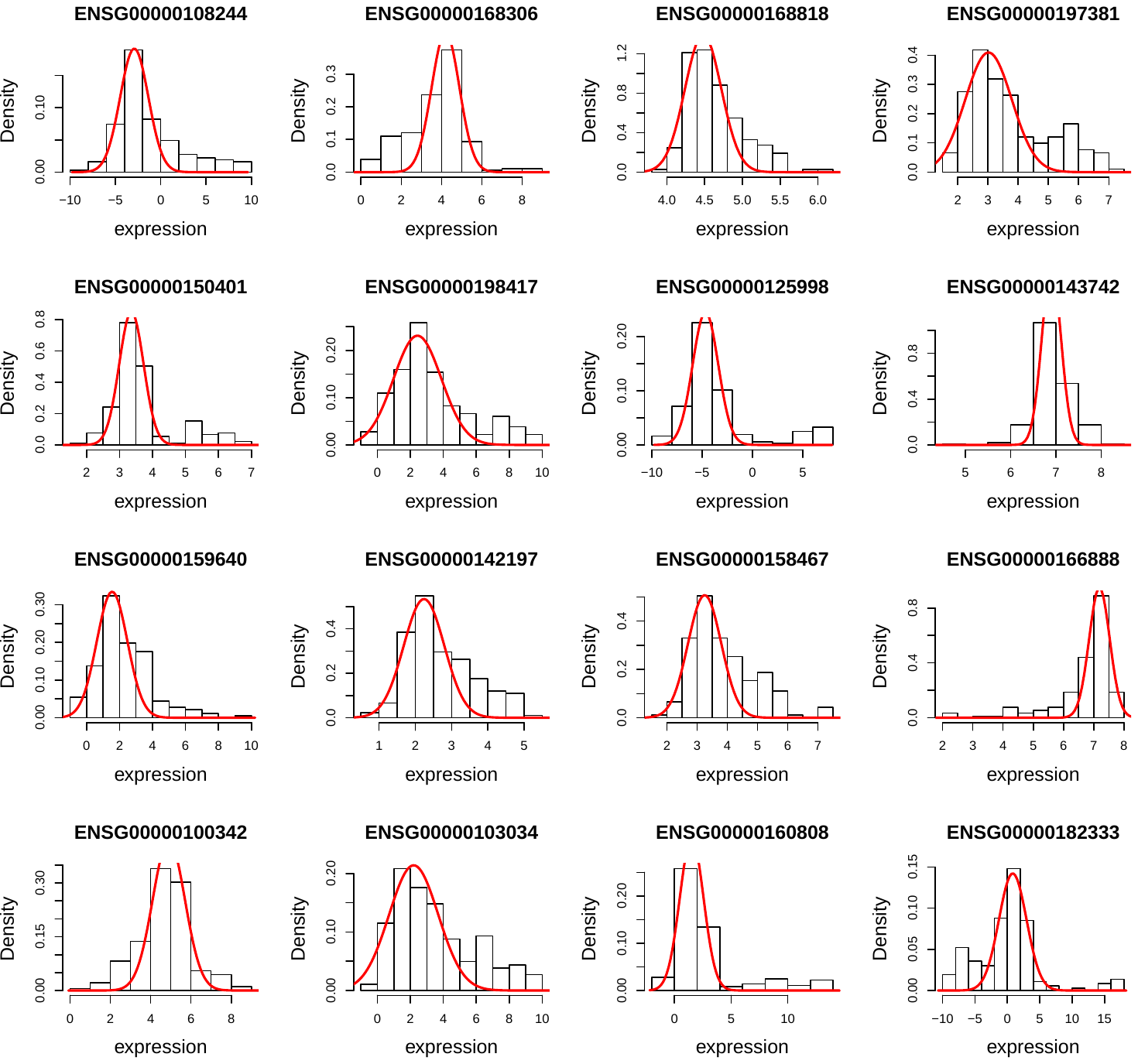

**Fig.S15. Histograms of RNA expressions in randomly selected genes whose ratio of and fitted standard deviation from AdaTiSS to MAD < 0.8.** The red curve is the density fitting from AdaTiSS.

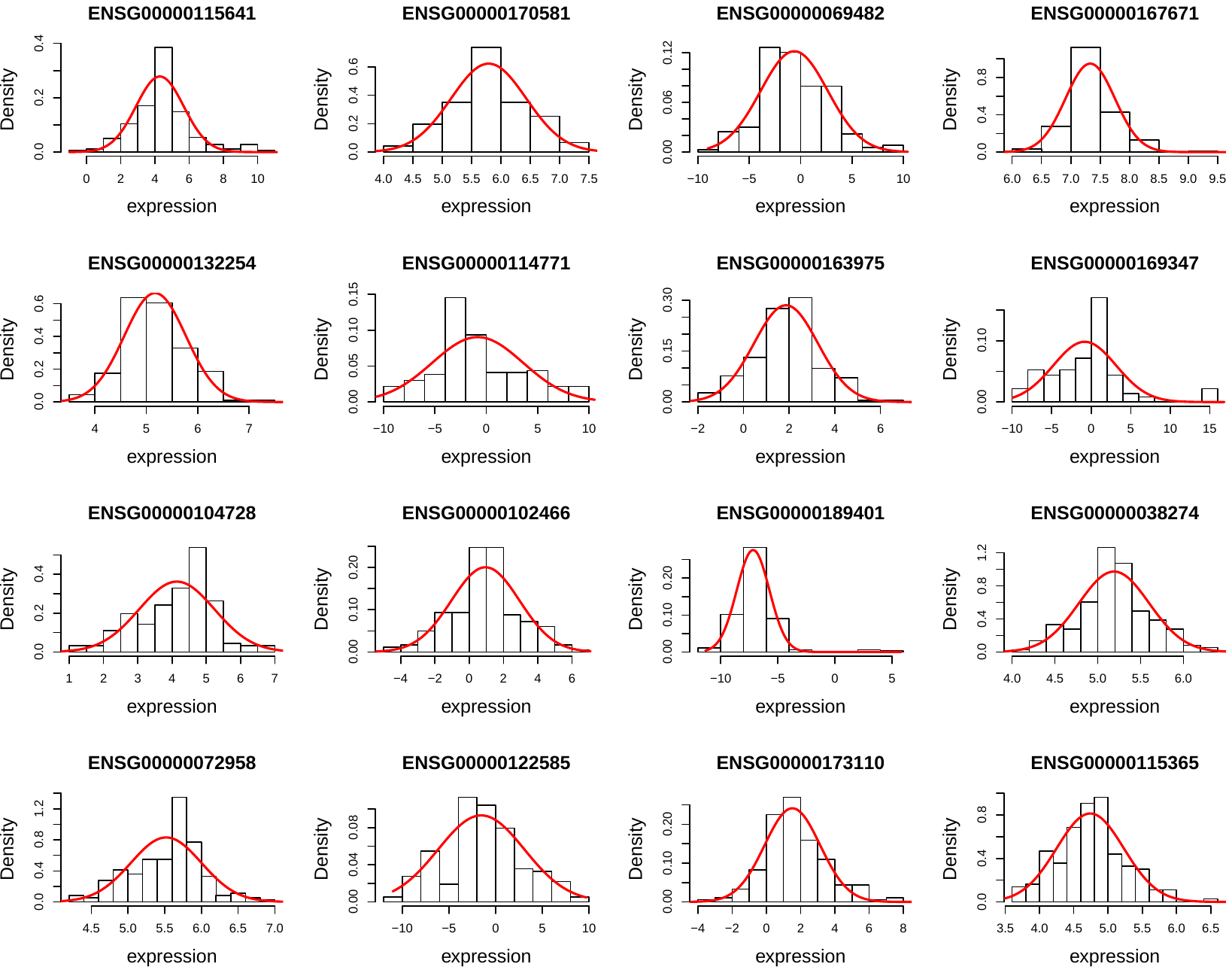

**Fig.S16. Histograms of RNA expressions in randomly selected genes whose ratio of and fitted standard deviation from AdaTiSS` to MAD > 1.2.** The red curve is the density fitting from AdaTiSS.

**Supplementary tables**

**Table S1. Gene information in the protein level.** The table provided identified protein list, raw protein abundances from 56 runs, normalized protein abundances, cleaned relative protein abundances in log scale, tissue medians from log of relative abundances, tissue standard z-scores based on median and MAD, and protein TS scores. For column name such as “i1r01s1.p05.Muscle Skeletal”, “i1” is for instrument id (i1, i2), “r01” for run id (r01, …, r28), “s1” for channel id (r1, r2 for 126, 131 reference channels, s1, .., s2 for tissue sample channels), “p05” for people id (p00 for reference), “Muscle Skeletal”‘ for tissue name (“ref” for reference samples). *Table is a .xlsx file.*

**Table S2. Gene information in the RNA level.** The table provided raw RNA TPM in log scale from 14 people, normalized RNA abundances, tissue medians from RNA abundances, tissue standard z-scores based on median and MAD, and RNA TS scores. *Table is a .xlsx file.*

**Table S3. Gene enrichment and concordance comparison between protein level and RNA.** The table provided protein and RNA combined information, and enrichment and concordance comparison in the gene level. *Table is a .xlsx file.*

**Table S4. GO and KEGG enrichment terms in protein.** The table provided the significant GO terms in biological process and significant KEGG pathways under FDR < 0.1 for the enriched proteins in individual tissues, and enriched GO terms for house-keeping proteins, and the significance (minus of log of p-values at base 10) for the significant metabolism pathways across tissues (at least having one p-value < 0.001 in one tissue). *Table is a .xlsx file.*

**Table S5. Isoform information in protein and RNA.** The table provided identified isoform names in protein and RNA, quantified protein isoform TS scores, RNA isoform TS scores from the corresponding genes (the genes without collapsed IDs defined in Section 2.3), protein and RNA enrichment comparison in isoforms, and tissue enriched protein isoform ranks in the corresponding RNA level in the same gene.

**Table S6. RNA and protein enrichment category counts comparison among protein-and-RNA commonly quantified genes.** “HK” stands for house-keeping, “ENCH not SPEC” for tissue-enriched-but-not-specific, “SPEC” for tissue-specific. The categories are defined in Supplementary Section 3.

| **Protein**  **RNA** | **HK** | **ENCH not SPEC** | **SPEC** | **other** | **Counts Sum** |
| --- | --- | --- | --- | --- | --- |
| **HK** | 622 | 647 | 123 | 1203 | 2595 |
| **ENCH not SPEC** | 477 | 2005 | 847 | 2007 | 5336 |
| **SPEC** | 75 | 278 | 332 | 412 | 1097 |
| **other** | 361 | 921 | 256 | 1672 | 3210 |
| **Counts Sum** | 1535 | 3851 | 1558 | 5294 | 12238 |

**Table S7. Protein identified genes across predicted protein class.**

| **Predicted protein class from HPA (total 19628)** | **Membrane**  **(total: 5455)** | **Intercellular**  **(total: 14889)** | **Secreted**  **(total: 2198)** |
| --- | --- | --- | --- |
| Identified genes in protein | 3143 | 10474 | 1984 |
| Unidentified genes in protein | 2312 | 4415 | 934 |
| p-value  From Fisher’s exact test | 0 | 1 | 1 |

**Table S8. Protein identified gene counts across RNA expression.**

| **RNA expression**  **(in log scale)** | $\boldsymbol{\leq1}$ | **(1,2]** | **(2,3]** | **(3,4]** | **(4,5]** |
| --- | --- | --- | --- | --- | --- |
| **protein identified**  **gene count** | 2199 | 1271 | 1708 | 2401 | 2336 |
| **protein unidentified**  **gene count** | 2756 | 772 | 624 | 450 | 310 |
| **RNA expression**  **(in log scale)** | **(5,6]** | **(6,7]** | **(7,8]** | **(8,9]** | **> 9** |
| **protein identified**  **gene count** | 1521 | 658 | 251 | 112 | 72 |
| **protein unidentified**  **gene count** | 157 | 64 | 12 | 9 | 7 |

**Table S9. Counts of identified and non-identified protein isoforms in rank 1 (a) and rank 2 (b) of RNA isoforms across RNA expression bins in log scale.**

| **(a) Counts of protein isoforms in rank 1 of RNA isoforms across RNA expression bins** | | | | | | |
| --- | --- | --- | --- | --- | --- | --- |
| **RNA expression (in log)** | $\boldsymbol{\leq1}$ | **(1,2]** | **(2,3]** | **(3,4]** | **(4,5]** | **> 5** |
| **not identified counts in protein** | 184 | 129 | 192 | 128 | 85 | 89 |
| **identified counts in protein** | 563 | 397 | 510 | 492 | 311 | 307 |
| chi-squared statistics = 10.285, degree of freedom = 5, p-value = 0.0675 from chi-square independence test | | | | | | |
| **(b) Counts of protein isoforms in rank 2 of RNA isoforms across RNA expression bins** | | | | | | |
| **RNA expression (in log)** | $\boldsymbol{\leq1}$ | **(1,2]** | **(2,3]** | **(3,4]** | **(4,5]** | **> 5** |
| **not identified counts in protein** | 2117 | 326 | 188 | 90 | 43 | 24 |
| **identified counts in protein** | 295 | 130 | 92 | 57 | 15 | 10 |
| chi-squared statistics = 181.06, degree of freedom = 5, p-value < $2.2\times{10}^{-16}$ from chi-square independence test | | | | | | |

**Table S10. Enrichment comparison in muscle gene groups.** The table provided protein and RNA enrichment comparison in gene and isoform levels in muscle gene groups. *Table is a .csv file.*

**Table S11. Tissue enriched proteins in genetic disease enrichment.** The table provided the gene names from the enriched proteins significantly enriched disease from OMIN database across tissues. *Table is a .csv file.*

**Table S12. Missing proteins.** The table provided the missing protein lists under different three different layers of criteria combined with protein and RNA enrichment information and HPA results. *Table is a .xlsx file.*

**Table S13. SNP variants.** The table provided the peptide sequences in SNP variants combined with outlier enrichment test results. *Table is a .csv file.*

**Table S14.** **Tissue full names and abbreviations.**

| **Tissue full name** | **Tissue name abbreviations** | **Tissue full name** | **Tissue name abbreviations** |
| --- | --- | --- | --- |
| Adrenal Gland | *Adrenal Gland* | Minor Salivary Gland | *Minor Salivary* |
| Artery - Aorta | *Artery Aorta* | Muscle - Skeletal | *Muscle Skeletal* |
| Artery - Coronary | *Artery Coronary* | Nerve - Tibial | *Nerve Tibial* |
| Artery - Tibial | *Artery Tibial* | Ovary | *Ovary* |
| Brain - Cerebellum | *Brain Cerebellum* | Pancreas | *Pancreas* |
| Brain - Cortex | *Brain Cortex* | Pituitary | *Pituitary* |
| Breast - Mammary Tissue | *Breast* | Prostate | *Prostate* |
| Colon - Sigmoid | *Colon Sigmoid* | Skin - Not Sun Exposed (Suprapubic) | *Skin Unexpo* |
| Colon - Transverse | *Colon Transverse* | Skin - Sun Exposed (Lower leg) | *Skin SunExpo* |
| Esophagus - Gastroesophageal Junction | *GE junction* | Small Intestine - Terminal Ileum | *Small Intestine* |
| Esophagus - Mucosa | *Esophagus Mucosa* | Spleen | *Spleen* |
| Esophagus - Muscularis | *Esophagus Muscle* | Stomach | *Stomach* |
| Heart - Atrial Appendage | *Heart Atrial* | Testis | *Testis* |
| Heart - Left Ventricle | *Heart Ventricle* | Thyroid | *Thyroid* |
| Liver | *Liver* | Uterus | *Uterus* |
| Lung | *Lung* | Vagina | *Vagina* |

**References**

1.	L. J. Carithers *et al.*, A novel approach to high-quality postmortem tissue procurement: the GTEx project. *Biopreservation and biobanking* **13**, 311-319 (2015).

2.	G. Consortium, The Genotype-Tissue Expression (GTEx) pilot analysis: multitissue gene regulation in humans. *Science* **348**, 648-660 (2015).

3.	N. A. Kulak, G. Pichler, I. Paron, N. Nagaraj, M. Mann, Minimal, encapsulated proteomic-sample processing applied to copy-number estimation in eukaryotic cells. *Nature methods* **11**, 319 (2014).

4.	J. Harrow *et al.*, GENCODE: the reference human genome annotation for The ENCODE Project. *Genome research* **22**, 1760-1774 (2012).

5.	M. Wang *et al.*, RobNorm: Model-Based Robust Normalization for High-Throughput Proteomics from Mass Spectrometry Platform. *bioRxiv*, 770115 (2019).

6.	M. Wang, L. Jiang, M. P. Snyder, AdaTiSS: A Novel Data-Adaptive Robust Method for Quantifying Tissue Specificity Scores. *(will appear in bioRxiv soon)*.

7.	L. v. d. Maaten, G. Hinton, Visualizing data using t-SNE. *Journal of machine learning research* **9**, 2579-2605 (2008).

8.	B. Efron. Local false discovery rates. (2005).

9.	H. Fujisawa, S. Eguchi, Robust parameter estimation with a small bias against heavy contamination. *Journal of Multivariate Analysis* **99**, 2053-2081 (2008).

10.	T. Kanamori, H. Fujisawa, Robust estimation under heavy contamination using unnormalized models. *Biometrika* **102**, 559-572 (2015).

11.	A. Ipsen, Derivation of the Statistical Distribution of the Mass Peak Centroids of Mass Spectrometers Employing Analog-to-Digital Converters and Electron Multipliers. *Analytical chemistry* **89**, 2232-2241 (2017).

12.	E. G. Hill *et al.*, A statistical model for iTRAQ data analysis. *Journal of proteome research* **7**, 3091-3101 (2008).

13.	M. Wang, L. Jiang, P. M. Snyder, AdaReg: Data Adaptive Robust Estimation in Linear Regression with Application in GTEx Gene Expressions. *(will appear in bioRxiv soon)*.

14.	M. Melé *et al.*, The human transcriptome across tissues and individuals. *Science* **348**, 660-665 (2015).

15.	X. Li *et al.*, The impact of rare variation on gene expression across tissues. *Nature genetics* **550**, 239 (2017).

16.	Y. Benjamini, Y. Hochberg, Controlling the false discovery rate: a practical and powerful approach to multiple testing. *Journal of the royal statistical society. Series B (Methodological)*, 289-300 (1995).

17.	G. O. Consortium, Gene Ontology Consortium: going forward. *Nucleic acids research* **43**, D1049–D1056 (2015).

18.	A. Franceschini *et al.*, STRING v9. 1: protein-protein interaction networks, with increased coverage and integration. *Nucleic acids research* **41**, D808-D815 (2012).

19.	M. Uhlén *et al.*, Tissue-based map of the human proteome. *Science* **347**, 1260419 (2015).

20.	J. S. Amberger, C. A. Bocchini, F. Schiettecatte, A. F. Scott, A. Hamosh, OMIM. org: Online Mendelian Inheritance in Man (OMIM®), an online catalog of human genes and genetic disorders. *Nucleic acids research* **43**, D789-D798 (2014).
